# Identification of parallel pH- and zeaxanthin-dependent quenching of excess energy in LHCSR3 in *Chlamydomonas reinhardtii*

**DOI:** 10.1101/2020.07.10.197483

**Authors:** Julianne M. Troiano, Federico Perozeni, Raymundo Moya, Luca Zuliani, Kwangryul Baek, EonSeon Jin, Stefano Cazzaniga, Matteo Ballottari, Gabriela S. Schlau-Cohen

**Author notes:** J.M.T. and F.P. contributed equally to this work. to whom correspondence should be addressed. (M.B.), (G.S.S.-C.).

## Abstract

Under high light conditions, oxygenic photosynthetic organisms avoid photodamage by thermally dissipating excess absorbed energy, which is called non-photochemical quenching (NPQ). In green algae, a chlorophyll and carotenoid-binding protein, light-harvesting complex stress-related (LHCSR3), detects excess energy via pH and serves as a quenching site. However, the mechanisms by which LHCSR3 functions have not been determined. Using a combined in vivo and in vitro approach, we identify two parallel yet distinct quenching processes, individually controlled by pH and carotenoid composition, and their likely molecular origin within LHCSR3 from *Chlamydomonas reinhardtii*. The pH-controlled quenching is removed within a mutant LHCSR3 that lacks the protonable residues responsible for sensing pH. Constitutive quenching in zeaxanthin-enriched systems demonstrates zeaxanthin-controlled quenching, which may be shared with other light-harvesting complexes. We show that both quenching processes prevent the formation of damaging reactive oxygen species, and thus provide distinct timescales and mechanisms of protection in a changing environment.

## Introduction

Sunlight is the essential source of energy for most photosynthetic organisms, yet sunlight in excess of the organism’s photosynthetic capacity can generate reactive oxygen species (ROS) that lead to cellular damage. To avoid damage, plants respond to high light by activating photophysical pathways that safely convert excess energy to heat, which is known as nonphotochemical quenching (NPQ) (Rochaix 2014). While NPQ allows for healthy growth, it also limits the overall photosynthetic efficiency under many conditions. If NPQ were optimized for biomass, yields would improve dramatically, potentially by up to 30% (Kromdijk et al. 2016, Zhu et al. 2010). However, critical information to guide optimization is still lacking, including the molecular origin of NPQ and the mechanism of regulation.

Green algae is a sustainable alternative for biofuels and livestock feed (Lum et al. 2013, Wijffels et al. 2010). In *Chlamydomonas (C.) reinhardtii*, the model organism for green algae, light-harvesting complex stress-related (LHCSR) is the key gene product for NPQ. LHCSR contains chlorophyll (Chl) and carotenoid (Car) held within its protein scaffold. Two isoforms of LHCSR, LHCSR1 and LHCSR3, are active in NPQ, although LHCSR3 is accumulated at higher levels and so has the major role (Dinc et al. 2016, Maruyama et al. 2014, Peers et al. 2009, Tokutsu et al. 2013). While the photophysical mechanism of quenching in light-harvesting complexes has not been determined, the primary proposals involve Chl-Car interactions (Liao et al. 2010, Ma et al. 2003, Ruban et al. 2007, Son, Pinnola, Gordon, et al. 2020, Son, Pinnola, and Schlau-Cohen 2020, de la Cruz Valbuena et al. 2019).

NPQ is triggered by a proton gradient across the thylakoid membrane that forms through a drop in luminal pH under excess light. The C-terminus of LHCSR3 contains a number of protonable residues exposed to the lumen that act as a pH sensor to trigger quenching (Ballottari et al. 2016, Liguori et al. 2013). The pH drop also activates the enzymatic conversion of the Car violaxanthin (Vio) to zeaxanthin (Zea) (Eskling et al. 1997). Along with LHCSR, other homologous light-harvesting complexes are likely involved in quenching (Nicol et al. 2019). In *C. reinhardtii*, the CP26 and CP29 subunits, which are minor antenna complexes of Photosystem II (PSII), have been implicated in NPQ (Cazzaniga et al. 2020). In higher plants, Zea has been reported to be involved in NPQ induction by driving light-harvesting complexes into a quenched state and/or by mediating interaction between light-harvesting complexes and PsbS, non-pigment binding subunits essential for NPQ induction in vascular plants (Sacharz et al. 2017, Ahn et al. 2008, Jahns et al. 2012). Similarly, Zea binding to LHCSR1 in the moss *Physcomitrella patens* and LHCX1 (a LHCSR homolog) in the microalga *Nannochloropsis oceanica* has been shown to be essential for NPQ (Pinnola et al. 2013, Park et al. 2019). Finally, in *C. reinhardtii*, a reduction of NPQ in the absence of Zea has been reported (Niyogi et al. 1997). In contrast, recent work has shown Zea to be unnecessary for NPQ induction or related to highly specific growth conditions (Bonente et al. 2011, Tian et al. 2019, Vidal-Meireles et al. 2020). Thus, the contribution of Zea to quenching in green algae remains under debate.

Because of the complexity of NPQ and the large number of homologous light-harvesting complexes, the individual contributions and mechanisms associated with LHCSR3, pH, and Zea have been challenging to disentangle, including whether they activate quenching separately or collectively. With the power of mutagenesis, the contribution of LHCSR3, and the dependence of this contribution on pH and Zea, can be determined. However, in vivo experiments leave the molecular mechanisms of LHCSR3 and its activation obscured. In vitro experiments, and particularly single-molecule fluorescence spectroscopy, are a powerful complement to identify protein conformational states (Gwizdala et al. 2016, Krüger et al. 2010, Kondo et al. 2017, Schlau-Cohen et al. 2014, Schlau-Cohen et al. 2015). A recent method to analyze single-molecule data, 2D fluorescence correlation analysis (2D-FLC) (Ishii et al. 2013a, Kondo et al. 2019), quantifies the number of conformational states and their dynamics, including simultaneous, independent processes. Thus, the conformational states associated with NPQ can be resolved.

Here, we apply a combined in vivo and in vitro approach to investigate NPQ in *C. reinhardtii*. We use mutagenesis, NPQ induction experiments, and fluorescence lifetime measurements on whole cells and single LHCSR3 complexes to show that pH and Zea function in parallel and can independently activate full quenching and prevent ROS accumulation. The pH-dependent quenching is controlled by protonable residues in the C-terminus of LHCSR3 as shown by mutagenesis to remove these residues. The Zea-dependent quenching is constitutive both in vitro and in vivo, reconciling previous conflicting reports. Based on the collective results, we find two likely quenching sites, *i.e.* Chl-Car pairs within LHCSR3, one regulated by pH and the other by Zea. The existence of two quenching processes provide different time scales of activation and deactivation of photoprotection, allowing survival under variable light conditions.

## Results

### Roles of pH and Zea in fluorescence intensity in vivo and in vitro

To disentangle the contributions of LHCSR, pH, and Zea, both in vivo and in vitro measurements were performed on different *C. reinhardtii* genotypes. Wild type (WT) strains (4A^+^ and CC4349), a strain depleted of LHCSR3 and LHCSR1 subunits (*npq4 lhcsr1*; Figure 1–– figure supplement 1 and 2)(Ballottari et al. 2016), a strain unable to accumulate Zea due to knock out of the enzyme responsible for xanthophyll cycle activation (*npq1*; Figure 1––figure supplement 3) (Li et al. 2016) and a mutant constitutively accumulating Zea due to knock out of the zeaxanthin epoxidase enzyme (*zep*; Figure 1––figure supplement 3) (Baek et al. 2016, Niyogi et al. 1997) were characterized in vivo. A mutant depleted of both LHCSR subunits (*npq4 lhcsr1*) was chosen rather than a mutant missing only LHCSR3 (*npq4*) due to the partial ability of LHCSR1 to substitute for LHCSR3 in its absence (Girolomoni et al. 2019).

To assess the ability of these phenotypes to undergo quenching of Chl fluorescence, the NPQ levels were measured in vivo after cells were acclimated to high light (HL, 500 μmol m^-2^s^-1^) for several generations to induce LHCSR expression (WT, *npq1*, and *zep* strains) and then exposed to strong light treatment (1500 μmol m^-2^s^-1^) for 60 minutes to induce maximum drop in luminal pH and Zea accumulation (WT, *np1, zep*, and *npq4 lhcsr1* strains; data for xanthophyll cycle activation shown in Figure 1––figure supplement 3). The NPQ induction kinetics are shown in Figure 1. In the WT strains, the maximum NPQ level was reached after 10 minutes of illumination and fully recovered in the dark (Figure 1A, black), despite a strong accumulation of Zea (Figure 1––figure supplement 3). In the *npq4 lhcsr1* strain, which lacks LHCSR subunits, a null NPQ phenotype was observed (Figure 1A, purple). These results confirm that LHCSR subunits are responsible for NPQ in *C. reinhardtii*.

**Figure 1.**
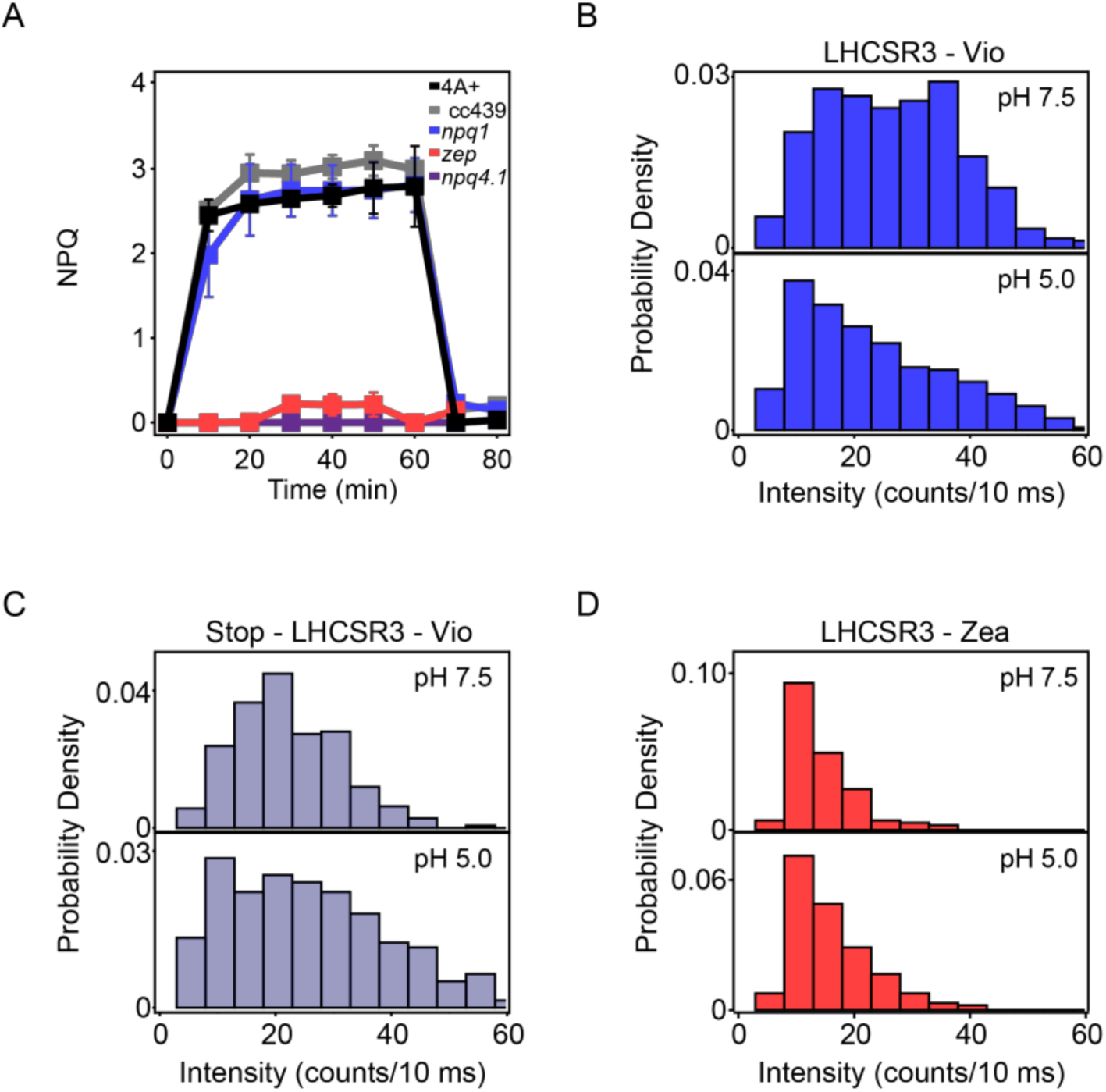
Experimental studies of quenching mechanisms in vivo and in vitro. (A) NPQ induction kinetics for high-light acclimated samples measured upon 60 minutes of high light (1500 μmol m^-2^s^-1^) in vivo. The results are reported as the mean of three independent biological replicates (N=3). Error bars are reported as standard deviation. Kinetics for low-light acclimated samples are shown in Figure 1 – figure supplement 5. Histograms of fluorescence intensities from single-molecule measurements on (B) LHCSR3-Vio, (C) stop-LHCSR3-Vio, and (D) LHCSR3-Zea at pH 7.5 (top) and pH 5.0 (bottom).

In the *npq1* strain, which lacks Zea, no reduction of the maximum level of NPQ was observed compared to its background, the 4A+ WT strain (Figure 1A, blue, black). The similar level and timescales of onset and recovery for NPQ suggest a minor role, if any, for Zea in light-activated quenching. In the *zep* strain, which constitutively accumulates Zea, a strong reduction of the NPQ level was observed compared to both WT strains CC4349 and 4A+ (Figure 1A, red). To understand why, first accumulation of the LHCSR subunits was measured (Figure 1––figure supplements 1 and 2). However, similar LHCSR3 content was found in WT strains (4A+ and CC4349), *npq1* and *zep* mutants. In the case of LHCSR1, similar accumulation was observed in 4A+ and *zep* mutant, while no trace of this subunit was detectable in the WT strain CC4349. Second, the extent of proton motive force as compared to WT was measured through the electrochromic shift (Bailleul et al. 2010). However, although proton transport into the lumen was reduced in the *zep* strain at low actinic light, it was similar at the higher irradiance used for measurement of NPQ (Figure 1––figure supplement 4). Therefore, neither differences in LHCSR accumulation nor in proton transport are the cause of the reduced NPQ phenotype in the *zep* mutant.

In order to investigate the effect of pH and Zea at the level of the LHCSR3 subunit, single-molecule fluorescence was measured for individual complexes (representative intensity traces in Figure 1––figure supplement 6). Histograms were constructed of the intensity levels for LHCSR3 with Vio at high and low pH, which mimic the cellular environment under low and high light conditions, respectively. The fluorescence intensity decreases, generally along with the fluorescence lifetime, as quenching increases. As an initial straightforward comparison, we present the fluorescence intensity from single LHCSR3. The fluorescence lifetime exhibits complex kinetics (de la Cruz Valbuena et al. 2019), and so we analyze the associated lifetime data with a recently-developed model-free method, 2D fluorescence lifetime correlation (2D-FLC), as discussed in detail below. As shown in Figure 1B, upon a decrease in pH from 7.5 to 5.0, the average fluorescence intensity of LHCSR3-Vio decreases as the quenched population increases, representing an increase in quenching of the excitation energy absorbed. This is consistent with the conclusion from the in vivo NPQ experiments that quenching can occur without Zea under high light conditions.

Activation of LHCSR3 as a quencher has been suggested previously to be related to protonation of putative pH-sensing residues present at the C-terminus (Figure 1––figure supplement 7). To assess the role of these protonable residues in pH-dependent quenching, a mutant of LHCSR3 lacking this protein portion (stop-LHCSR3) was measured in vitro (Figure 1––figure supplements 8 and 9). Upon the same pH decrease that induced quenching in LHCSR3-Vio, stop-LHCSR3-Vio exhibits similar fluorescence intensity (Figure 1C). The data show that the mutants have lost the ability to activate quenching channels upon a pH drop, highlighting the sensing role of the protonable residues of the C-terminus of LHCSR3.

Single-molecule measurements were also performed on LHCSR3 with Zea at high and low pH. Under both conditions, as shown in Figure 1D, LHCSR3 with Zea in vitro show decreased fluorescence intensity due to increased quenching. The pH-independence of these histograms is consistent with the NPQ measurements in the *zep* mutants where high light, and the associated pH drop in the lumen, does not change quenching levels. However, these measurements point to the existence of a constitutive quenching process in the presence of Zea, consistent with in vivo fluorescence lifetime measurements discussed below.

### Roles of pH and Zea in fluorescence lifetime in vitro

To further assess and quantify the contributions of pH and Zea in quenching in LHCSR3 in vitro, we used 2D-FLC to analyze the fluorescence lifetime data from single LHCSR3 complexes (Kondo et al. 2019). The 2D-FLC analysis identifies fluorescence lifetime states, which correspond to protein conformations with different extents of quenching, and quantifies the transition rates between the lifetime states, which correspond to switches between the protein conformations. The analysis also identifies the number of dynamic components that switch between the states, which correspond to independent processes occurring simultaneously yet separately within single LHCSR3.

The number of lifetime states was determined through the lifetime distribution (Figure 2 ––figure supplement 1 and 2). The distribution was constructed by performing an inverse Laplace transform of the lifetime data (time between excitation and emission), which was recorded on a photon-by-photon basis. Unlike traditional lifetime fitting, the distribution is a model-free analysis of the decay components, which is particularly important when there are multiple and varied components as is the case for LHCSR3 (de la Cruz Valbuena et al. 2019). This allows analysis of multi-component lifetimes, even for the low signal-to-background regime of single-molecule data. For each of the LHCSR3 samples, two lifetime states were observed, an active state (∼2.5 ns) and a quenched state (∼0.5 ns).

**Figure 2.**
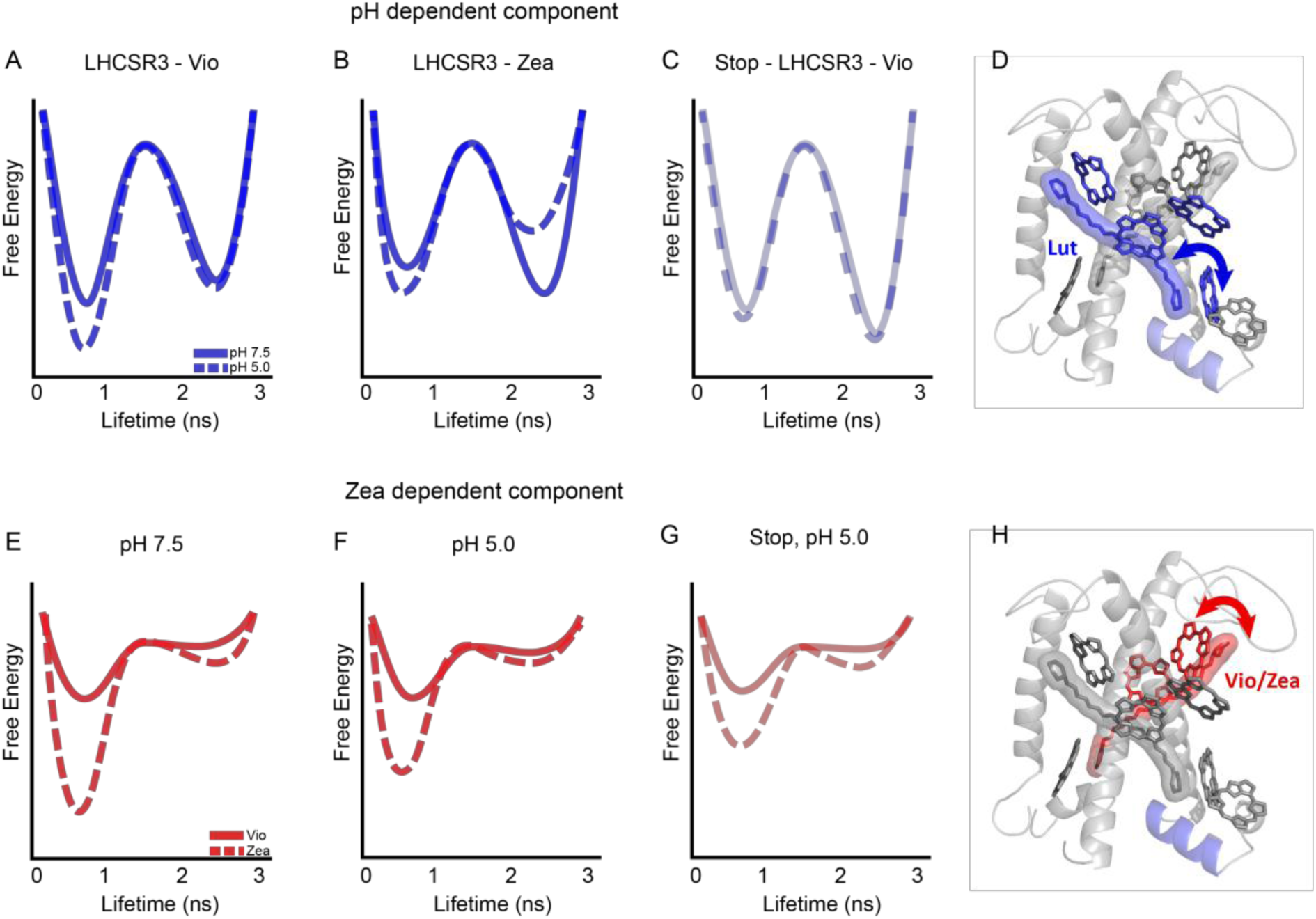
pH- and Zea-dependent quenching in LHCSR3. Free energy diagrams extracted from the 2D-FLC analysis (A-C and E-G) and structural model with likely quenching sites (D and H) for the effects of pH (top) and Zea (bottom) on protein dynamics.

The number of parallel conformational processes occurring within single LHCSR3 was found by quantifying the number of dynamical components through a global fit of the auto- and cross-correlations for each sample. The best fits to the data were accomplished by including three dynamical components at different timescales (Figure 2––figure supplement 3, Figure 2 – table supplement 1). Two of these components, one on a tens of microsecond timescale and the other on a hundreds of microsecond timescale, would be hidden in traditional single-molecule analyses. The cross-correlation for every LHCSR3 sample begins above zero (Figure 2––figure supplement 3), which appears when the dynamic components occur in parallel (Kondo et al. 2019). The Chl *a* have the lowest energy levels, which are the emissive states that give rise to each component. The existence of three components indicates multiple Chl *a* emissive sites within LHCSR3, consistent with previous models of LHCs (Mascoli et al. 2019, Mascoli et al. 2020, Krüger et al. 2010, Krüger et al. 2011). The two dynamic components arise from changes in the extent of quenching of the Chl emitters. The third component is static at <0.01 s, which is attributed to unquenched emitters in the active state and partially photobleached complexes in the quenched state.

The parameters extracted from the global fit include the intensity of and population in each lifetime state. The rate constants for the transitions between the states are also determined, primarily from the cross-correlation functions. The parameters can be used to construct illustrative free energy landscapes, which are shown in Figure 2 for the two dynamic components. The depth of each free energy well depends on the population of the state and the barrier between wells depends on the rate constants (Kondo et al. 2019). The energies associated with each well are given in Figure 2––table supplement 1.

We examine the dependence on molecular parameters of the two dynamical components. Figure 2A and 2B show the pH-dependence of the free energy landscapes for the slower component in LHCSR3-Vio and LHCSR3-Zea, respectively. In both cases, a decrease in pH from 7.5 to 5.0 stabilizes the quenched state. In LHCSR3-Vio, the quenched state is stabilized by a decrease in the transition rate from the quenched to the active state, corresponding to a higher barrier in the free energy landscape. In LHCSR3-Zea, the decrease in the transition rate from the quenched to the active state is also accompanied by an increase in the transition rate from the active to the quenched state, further stabilizing the quenched state relative to the unquenched one. In stop-LHCSR3-Vio, however, no change in the population of the quenched state is observed upon a decrease in pH (Figure 2C), reflecting the expected pH-independence of the sample.

Figure 2E and F show the Zea-dependence of the free energy landscapes of LHCSR3 for the faster dynamic component at pH 5.0 and pH 7.5. At both pH levels, conversion from Vio to Zea stabilizes the quenched state via a decrease in the transition rate from the quenched to active state. At pH 5.0, the transition rate to the quenched state increases as illustrated by the lower barrier, which would enable rapid equilibration of population into the quenched state. The Zea-dependent behavior is maintained for stop-LHCSR3 (Figure 2G), where the quenched state is still stabilized in the presence of Zea.

The two dynamic components reveal two parallel yet independent photoprotective processes, one pH-dependent and one Zea-dependent, within LHCSR3, demonstrating multifunctionality of the protein structure. Each component likely arises from a Chl-Car pair, where the Car can quench the emissive Chl. The two components are both biased towards the quenched state in the LHCSR3 complexes, consistent with previous work where a quenching component was found to be active in LHCSR3, even at pH 7.5 (de la Cruz Valbuena et al. 2019). By considering the results of the 2D-FLC analysis along with previous structural, spectroscopic and theoretical work, we speculate as to the likely sites associated with each component. Although no high-resolution structure of LHCSR3 has been determined, we illustrate possible quenching sites (Figure 2D and H) within a working structural model of LHCSR3 (Bonente et al. 2011). As shown in Figure 2D and H (Figure 2––table supplement 2), LHCSR3 purified from *C. reinhardtii* contains eight Chl molecules (7-8 Chl *a* and 0-1 Chl *b* molecules) and two Cars (one lutein (Lut) and one Vio or Zea). Based on sequence comparison with LHCII and CP29, the conserved Chl *a* binding sites are the following: Chls *a* 602, 603, 609, 610, 612 and 613, with Chls *a*604, 608 and 611 proposed as well (Bonente et al. 2011, Liguori et al. 2016). Previous spectroscopic analysis of LHCSR3 from *C. reinhardtii* has identified the likely binding sites of Lut and Vio/Zea within the structural model (Bonente et al. 2011). Given that there are two Cars bound at the internal sites of LHCSR3, it is likely that each Car and its neighboring Chl is the major contributor for one of the two dynamic components.

The pH-dependent component (Figure 2, top, blue) likely involves Lut and its neighboring Chl. Both Chl *a* 612 (coupled to Chl *a*s 610 and 611) and Chl *a* 613 have previously been proposed as quenching sites given their physical proximity to the Lut (Liguori et al. 2016, Ruban et al. 2007). The Chl *a* 610-612 site contains the lowest energy Chl *a*, which has been shown to be a major energy sink and thus the primary emitter (Muh et al. 2010, Schlau-Cohen et al. 2009, Novoderezhkin et al. 2011). Additionally, computational results have shown that the interaction between the Lut site and Chl *a* 612 has large fluctuations (Liguori et al. 2015). This agrees with the slower dynamics found for this component. However, recent in vivo and in vitro analyses found that the removal of Chl *a* 613 results in a reduction in LHCSR specific quenching, while removal of Chl *a* 612 only affected which Chl was the final emitter of the complex (Perozeni et al. 2019). While either of these sites are potential quenching sites, it is likely that Chl *a* 613 plays the major role in pH-dependent quenching in LHCSR3 in *C. reinhardtii*.

With a decrease in pH from 7.5 to 5.0, the equilibrium free energy difference for the pH-dependent component is shifted toward the quenched state by over 200 cm^-1^ in LHCSR3-Vio and over 500 cm^-1^ in LHCSR3-Zea. The specific conformational change upon protonation that leads to this stabilization remains undetermined. However, proposals in the literature include reduced electrostatic repulsion in the lumen-exposed domain causes a change in the distance and/or orientation between the helices (Ballottari et al. 2016) and an increase in protein-protein interactions (de la Cruz Valbuena et al. 2019). These conformational changes could produce a displacement of Lut towards Chl *a* 613. The negligible (< 30 cm^-1^) change in the equilibrium free energy difference for stop-LHCSR3 (Figure 2C, Figure 2––table supplement 1) demonstrates the functional role of the C-terminal tail in the conformational change.

The Zea-dependent component (Figure 2, bottom, red) likely involves Vio/Zea and the neighboring Chl *a*s 602-603 (Bonente et al. 2011, Di Valentin et al. 2009, Lampoura et al. 2002). With conversion from Vio to Zea, the free energy landscape changes significantly, and thus is likely to involve Vio/Zea itself. In addition, MD simulations have shown this Car site to be highly flexible, sampling many configurations (Liguori et al. 2017), which is consistent with the faster dynamics observed here. Upon substitution of Zea for Vio, the equilibrium free energy difference becomes further quenched-biased by over 550 cm^-1^ at pH 7.5 and over 300 cm^-1^ at pH 5.0. This result is consistent with a role of Zea in quenching of LHCSR3 that does not require a decrease in pH and therefore being unrelated to the light dependent NPQ observed in vivo in the WT that almost completely recovered in the dark (Figure 1A).

In the stop-LHCSR3, the equilibrium free energy differences for the Zea-dependent (faster) component is similar to the wild type samples (Figure 2G). This is consistent with the Vio/Zea-Chl *a* 602-603 site as the major contributor for this component. Although qualitatively similar, there is a small decrease (<200 cm^-1^) in the stabilization of the quenched state upon Zea incorporation. This difference suggests that the C-terminal tail has an allosteric effect throughout the protein.

The static component, which is assigned to unquenched emitters in the active state and partially photobleached ones in the quenched state, has a large contribution to the correlation profiles (Figure 2––table supplement 1). The large amplitude is consistent with the low number of Cars available to interact with the Chls and thus the presence of several unquenched emissive Chl *a*. Given the structural arrangement of the Cars and Chls, the static component is likely due to Chls 604, 608 and 609, which sit far from the Cars.

### Roles of pH and Zea in fluorescence lifetime in vivo

Quenching mechanisms were further investigated in vivo by measuring fluorescence emission lifetimes of whole cells acclimated to low or high light at 77K, as traditional NPQ measurements can be affected by artifacts (Tietz et al. 2017). Under these conditions, the photochemical activity is blocked and by following the emission at 680 nm it is possible to specifically investigate the kinetics of PSII, the main target of NPQ. Cells were either grown in low or high light, which determines the level of LHCSR protein (Figure 1––figure supplement 1 and 2) and the fluorescence lifetime was recorded before and after exposure to 60 minutes of high light (1500 μE), which induces ΔpH and determines the level of Zea. These light conditions, combined with the genotypes generated, enabled studies that partially or fully separated the contributions of the different components of NPQ.

Whole cell fluorescence lifetime traces show that LHCSR is necessary for NPQ in *C. reinhardtii*. WT and *npq1*, which lacks Zea, cells grown in high light show a faster fluorescence decay, or an increase in quenching, after exposure to 60 minutes of high light (Figure 3A, gray bars, fluorescence decays and fits to data shown in SI). For *npq4 lhcsr1* cells, which lack LHCSR, similar fluorescence decay kinetics were measured regardless of light treatment (Figure 3A, purple), which is comparable to the unquenched kinetics of WT cells. WT and *npq1* cells grown in control light (low LHCSR content) remain unquenched, even after exposure to 60 minutes of high light (Figure 3––figure supplement 1 and 2). These results are consistent with the NPQ measurements shown in Figure 1A. Similar to WT, *npq1* cells grown in high light show a faster fluorescence decay after exposure to 60 minutes of high light (Figure 3A, blue bars). While the results from WT show a role for pH and/or Zea in light-induced quenching in LHCSR3, the results from the *npq1* strain show that quenching can occur without Zea, *i.e.*, induced by the pH drop alone.

**Figure 3.**
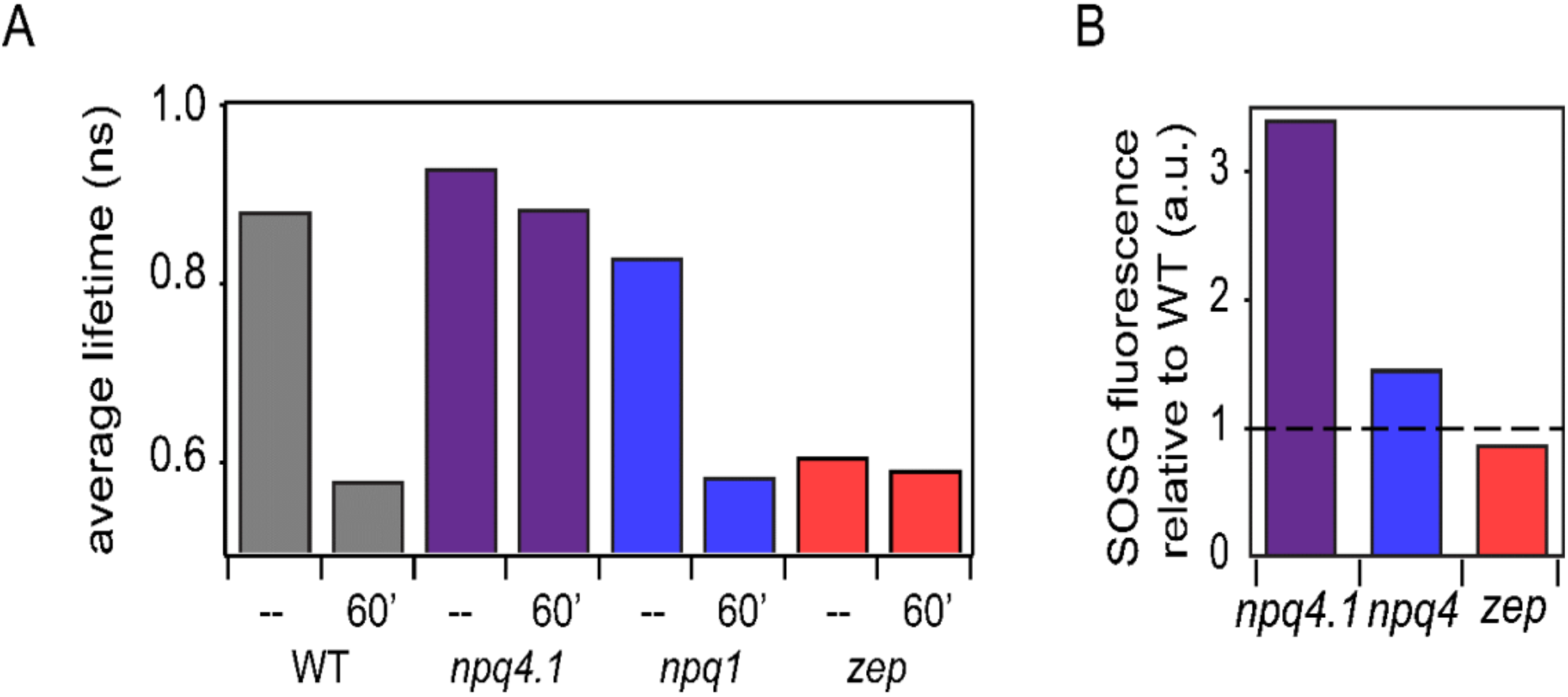
Fluorescence lifetime decay of *Chlamydomonas reinhardtii* whole cells at 77K and singlet oxygen formation. (A) Fluorescence lifetimes were measured on high light (600 μmol m^-2^s^-1^) acclimated samples. Each genotype was measured at a dark-adapted state (--) or after 60 minutes of high light treatment (2000 μmol m^-2^s^-1^, 60’). WT samples shown here are 4A^+^ strain. Similar results were obtained in the case of CC4349 strain. The *npq4 lhcsr1* mutant is indicated here as *npq4.1*. Fluorescence lifetime traces for all genotypes and light conditions are shown in Figure 3––figure supplement 1 and 2 with fit values in Figure 3 – table supplement 1. (B) Singlet oxygen production rates for high light acclimated samples relative to WT (4A+ for *npq1* and *npq4 lhcsr1*, CC4349 for *zep*). Dotted line represents WT level at 1. The results reported are representative of three independent biological replicates for each genotype in LL or HL. Original data are reported in figure supplement 3. Singlet oxygen kinetics are shown in Figure 3 – figure supplement 3. Low light acclimated samples are shown in Figure 3––figure supplement 4.

The *zep* mutant presented a similar decay among all samples, regardless of high or low light acclimation or light treatment, that was much faster, or more quenched, compared to the decay of dark-adapted WT (Figure 3A, red, Figure 3––figure supplement 1 and 2). These results indicate a constitutive quenching upon Zea accumulation, consistent with the pH-independent quenching observed in the results in Figure 1A and 1D and Figure 2E and 2F. However, the constitutive quenching observed in the *zep* mutant was unchanged in low vs. high light acclimated *zep* cells suggesting that the Zea dependent quenching observed in *zep* mutants is a more general process as opposed to one that occurs solely in LHCSR3.

### Role of Zea and NPQ in photoprotection

The main function of quenching the Chl singlet excited states is to alleviate the excitation energy pressure on the photosynthetic apparatus, thereby preventing ROS formation and subsequent photoinhibition of their primary target, PSII. Singlet oxygen is the main ROS species formed at the level of PSII. In order to correlate the NPQ levels and quenching measured with ROS formation, singlet oxygen production was recorded in the different genotypes herein investigated (Figure 3B, Figure 3––figure supplement 3 and 4). As expected from the low level of NPQ induction, *npq4* lhcsr1, which lacks LHCSR, demonstrated the highest level of singlet oxygen production, regardless of light treatment. Interestingly, the effect of Zea was almost negligible in high light acclimated samples (with a very high NPQ induction). Notably, the amount of singlet oxygen production was correlated with average lifetime (Figure 3A), *i.e.*, inversely correlated with quenching, confirming that the quenching of Chl singlet excited states investigated here plays a role in photoprotection.

## Discussion

This work leverages in vivo and in vitro experimental approaches to investigate NPQ mechanisms in *C. reinhardtii* and the molecular parameters responsible for their activation. In higher plants, both lumen acidification and Zea accumulation have been long understood to play a role in the induction of NPQ. While lumen acidification was thought to play a similar role in *C. reinhardtii*, here we characterize the impact of Zea accumulation, which had previously been elusive. We further identify the likely the molecular mechanisms of both pH and Car composition.

Both our in vivo and in vitro results point to pH and Zea controlling separate quenching processes that independently provide photoprotection. Full light-induced quenching upon lumen acidification in the *npq*1 strain, which lacks Zea, and the full constitutive quenching in the *zep* strain, which is Zea-enriched, demonstrate two separate quenching and induction processes in vivo. Likewise, the 2D-FLC analysis on single LHCSR3 quantified two parallel dynamic components, or separate quenching processes, one of which is pH-dependent and the other Zea-dependent.

For pH-dependent quenching, our 2D-FLC results, along with previous mutagenesis, (Perozeni et al. 2019) point to Lut-Chl613 as the likely quenching site. Analysis of stop-LHCSR3, which lack the protonable residues in the C terminus, definitively shows that the C terminus controls quenching activity by pH-induced stabilization of the quenched conformation of LHCSR3. The 2D-FLC results show that removal of the C-terminal tail removes pH-dependent quenching, while leaving Zea-dependent quenching nearly unaffected. Analogously, the WT low light grown strains, which lack LHCSR, also lack the ability for NPQ induction, supporting the critical role of the protonable residues unique to LHCSR in activating quenching.

For Zea-dependent quenching, we observe a constitutive quenching process both in vivo and in vitro that we assign to the Vio/Zea-Chl *a*602-603 site. Constitutive, as opposed to the typical inductive quenching, represents a distinct timescale and mechanism for the effect of Zea. The Zea-dependent quenching mechanism is described at the molecular level in the case of LHCSR3 but it likely shared with our LHC proteins: indeed, a strong reduction of fluorescence lifetime can be observed in whole cells in the case of *zep* mutant, even in low light acclimated cells where the amount of LHCSR3 is minimal (Figure 3––figure supplement 1 and 2). This demonstrates that LHCSR subunits are not the sole quenching site where Zea-dependent quenching occurs, consistent with previous work implicating the minor antenna complexes (Cazzaniga et al. 2020). It is important to note that in the case of *zep* mutant, not only does Zea completely substitute Vio (de-epoxidation index is 1, Figure 1 –– figure supplement 3), but also the Zea/Chl ratio is much higher compared to the ratio observed in WT or *npq4 lhcsr1*. This suggests an alternative possibility where the strong quenching observed in the *zep* mutant could be related to accumulation of Zea in the thylakoid membrane changing the environment where the photosystems and light-harvesting complexes are embedded, inducing the latter to a strong quenched state. While both possibilities allow for constitutive quenching in the presence of Zea, it is the constitutive quenching itself that is potentially the origin of the conflicting results in the literature.

Taken together, the in vivo and in vitro results demonstrate that separate pH- and Zea-dependent quenching processes exist and both provide efficient photoprotection. While in vivo measurements suggest that pH-dependent quenching is often dominant over Zea-dependent quenching, and correspondingly more efficient in photoprotection, the conformational states and pigment pairs likely responsible function in parallel via similar mechanisms. However, the timescales and induction associated with each quenching process are distinct; responsive pH-dependent quenching works in combination with the guaranteed safety valve of Zea-dependent quenching, potentially to protect against a rapid return to high light conditions.

## Materials and Methods

### Strains and culture conditions

*C. reinhardtii* WT (4A^+^and CC4349) and mutant strains (*npq1, npq4 lhcsr1* in 4A^+^background, *zep* in CC4349 background) were grown at 24°C under continuous illumination with white LED light at 80 μmol photons m^-2^ s^-1^ (low light, LL) in high salts (HS) medium (Harris et al. 2008) on a rotary shaker in Erlenmeyer flasks. High light (HL) acclimation was induced by growing cells for 2 weeks at 500 μmol photons m^-2^ s^-1^ in HS.

### SDS-PAGE Electrophoresis and Immunoblotting

SDS–PAGE analysis was performed using the Tris-Tricine buffer system (Schägger et al. 1987). Immunoblotting analysis was performed using αCP43 (AS11 1787), αPSAA (AS06 172) and αLHCSR3 (AS14 2766) antibodies purchased from Agrisera (Sweden).

### Violaxanthin de-epoxidation kinetics and pigment analysis

Violaxanthin de-epoxidation kinetics were performed by illuminating the different genotypes with red light at 1500 μmol photons m^-2^ s^-1^ up to 60 minutes. Pigments were extracted 80% acetone and analysed by HPLC as described in (Lagarde et al. 2000).

### NPQ and electrochromic shift measurements

NPQ induction curves were measured on 60 minutes dark adapted intact cells with a DUAL-PAM-100 fluorimeter (Heinz-Walz) at room temperature in a 1×1 cm cuvette mixed by magnetic stirring. Red saturating light of 4000 μmol photons m^-2^ s^-1^ and red actinic light of 1500 μmol photons m^-2^ s^-1^ were respectively used to measure Fm and Fm’ at the different time points. NPQ was then calculated as Fm/Fm’-1. Proton motive force upon exposure to different light intensities was measured by Electrochromic Shift (ECS) with MultispeQ v2.0 (PhotosynQ) according to Kuhlgert, S. et al. MultispeQ Beta: A tool for large-scale plant phenotyping connected to the open photosynQ network (Kuhlgert et al. 2016).

### LHCSR3 WT and Stop proteins refolding for in vitro analysis

pETmHis containing LHCSR3 CDS previously cloned as reported in Perozeni et al. 2019 served as template to produce Stop mutant using Agilent QuikChange Lightning Site-Directed Mutagenesis Kit. Primer TGGCTCTGCGCTTCTAGAAGGAGGCCATTCT and primer GAATGGCCTCCTTCTAGAAGCGCAGAGCCA were used to insert a premature stop codon to replace residue E231, generating a protein lacking 13 c-terminal residues. LHCSR3 WT and Stop protein were overexpressed in BL21 *E. coli* and refolded *in vitro* in presence of pigments as previously reported (Bonente et al. 2011). Pigments used for refolding were extracted from spinach thylakoids. Vio or Zea-binding versions of LHCSR3 were obtained using Vio or Zea containing pigment extracts in the refolding procedure. Zea-containing pigments were obtained by *in vitro* de-epoxidation (de la Cruz Valbuena et al. 2019, Pinnola et al. 2017) Fluorescence emission at 300K with excitation at 440 nm, 475 nm and 500 nm was used to evaluate correct folding as previously reported (Ballottari et al. 2010).

### Single-molecule fluorescence spectroscopy

12 μM solutions of purified LHCSR3 complexes were stored at −80°C. Immediately prior to experiments, LHCSR3 samples were thawed over ice and diluted to 50 pM using buffer containing 0.05% n-dodecyl-α-D-maltoside and either 20 mM HEPES-KOH (pH 7.5) or 40 mM MES-NaOH (pH 5.0). The following enzymatic oxygen-scavenging systems were also used: (1) 25 nM protocatechuate-3,4-dioxygenase and 2.5 mM protocatechuic acid for pH 7.5 and (2) 50 nM pyranose oxidase, 100 nM catalase and 5 mM glucose for pH 5.0.(Aitken et al. 2008, Swoboda et al. 2012) Diluted solutions were incubated for 15 minutes on Ni-NTA coated coverslips (Ni_01, Microsurfaces) fitted with a Viton spacer to allow LHCSR3 complexes to attach to the surface via their His-tag. The sample was rinsed several times to remove unbound complexes and sealed with another coverslip.

Single-molecule fluorescence measurements were performed in a home-built confocal microscope. A fiber laser (FemtoFiber pro, Toptica; 130 fs pulse duration, 80 MHz repetition rate) was tuned to 610 nm and set to an excitation power of 350 nW (2560 nJ/cm^2^ at the sample plane, assuming a Gaussian beam). Sample excitation and fluorescence collection were accomplished by the same oil-immersion objective (UPLSAPO100XO, Olympus, NA 1.4). The fluorescence signal was isolated using two bandpass filters (ET690/120x and ET700/75m, Chroma). The signal was detected using an avalanche photodiode (SPCM-AQRH-15, Excelitas) and photon arrival times were recorded using a time-correlated single photon counting module (Picoharp 300, Picoquant). The instrument response function was measured from scattered light to be 380 ps (full width at half maximum). Fluorescence intensity was analyzed as described previously using a change-point-finding algorithm (Watkins et al. 2005). Fluorescence emission was recorded until photobleaching for the following number of LHCSR3 in each sample: 132 LHCSR3-Vio at pH 7.5 (1.6•10^7^ photons); 173 LHCSR3-Vio at pH 5.5 (1.3•10^7^ photons); 95 LHCSR3-Zea at pH 7.5 (1.4•10^7^ photons); 216 LHCSR3-Zea at pH 5.5 (9.0•10^6^ photons); 125 stop-LHCSR3-Vio at pH 7.5 (2.5•10^7^ photons); 130 stop-LHCSR3-Vio at pH 5.5 (7.9•10^6^ photons); 148 stop-LHCSR3-Zea at pH 7.5 (1.3•10^7^ photons); 116 stop-LHCSR3-Zea at pH 5.5 (9.9•10^6^ photons). Each data set was collected over two or three days for technical replicates and the distribution generated each day was evaluated for consistency.

### 2D Fluorescence lifetime correlation analysis

2D fluorescence lifetime correlation analysis was performed as detailed previously (Kondo et al. 2019). Briefly, we performed the following analysis. First, the total number of states exhibiting distinct fluorescence lifetimes was estimated from the 1D lifetime distribution. The lifetime distribution is determined using the maximum entropy method (MEM) to perform a 1D inverse Laplace transform (1D-ILT) of the 1D fluorescence lifetime decay (Ishii et al. 2013a). Next, a 2D fluorescence decay (2D-FD) matrix was constructed by plotting pairs of photons separated by ΔT values ranging from 10^−4^ to 10 seconds. The 2D-FD matrix was transformed from t-space to the 2D fluorescence lifetime correlation (2D-FLC) matrix in τ-space using a 2D-ILT by MEM fitting (Ishii et al. 2012, 2013a, b). The 2D-FLC matrix is made up of two functions: the fluorescence lifetime distribution, A, and the correlation function, G. In practice, the initial fluorescence lifetime distribution, A_0_, was estimated from the 2D-MEM fitting of the 2D-FD at the shortest ΔT (10^−4^ s). Then the correlation matrix, G_0_, was estimated at all ΔT values with A_0_ as a constant. A_0_ and G_0_, along with the state lifetime values determined from the 1D analysis, were used as initial parameters for the global fitting of the 2D-FDs at all ΔT values. A was treated as a global variable and G was treated as a local variable at each ΔT (now G(ΔT)). The resulting fit provides the correlation function, G(ΔT). The correlation function was normalized with respect to the total photon number in each state. Each set of correlation curves (auto- and cross-correlation for one sample) were globally fit using the following model function:

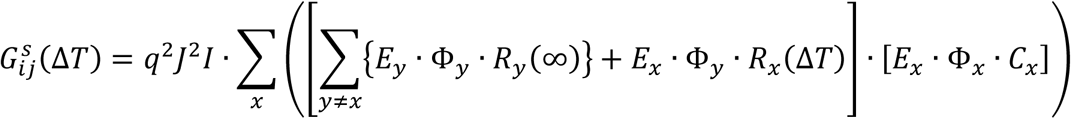

This equation accounts for multiple, independent emitters within one protein (multiple components). Here, x and y indicate the component number, i and j indicate the state (auto correlation for i=j, cross correlation for i ≠ j), q accounts for experimental factors such as the detection efficiency, filter transmittance, gain of the detector, etc., J is the laser power, and I is the total photon number proportional to the total measurement time. E, Φ, and C are diagonal matrices composed of the optical extinction coefficient, fluorescence quantum yield, and state population, respectively. R is a matrix element that is related to the rate matrix.

The rate constants determined from the 2D-FLC analysis were used to calculate the free energies for each protein state shown in Figure 2A-C and 2E-G. The rate constants for transitions between the quenched and active states are related to the free energies associated with both states through the Arrhenius equation:

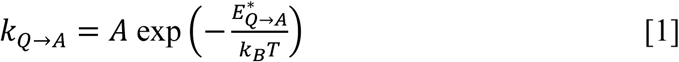

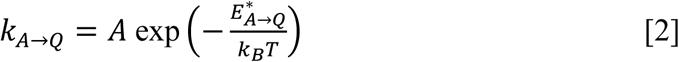

Here, *k*_*Q*→*A*_ and 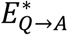 (*k*_*A*→*Q*_ and 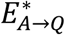) are the rate constant and activation energy, respectively, for the transition from the quenched (Q) to the active (A) state. A is a constant, *k*_*B*_ is the Boltzmann constant, and T is the temperature. Upon equilibration of the Q and A states, the free-energy difference, Δ*E*^∗^, is given by the following equation:

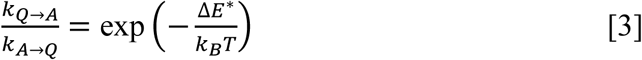

Using the dynamic rates determined from the fits to the correlation function, we calculated Δ*E*^∗^ at T = 300 K. The free-energy differences between the quenched and active states are shown as the energetic differences between the minima in the energy landscapes shown in Figure 2. The potential barriers were scaled by assuming the constant A in Equations [1] and [2] to be 1000, which was shown previously to be a reasonable estimate for LHCSR1 (Kondo et al. 2019).

### 77K fluorescence

Low temperature quenching measures were performed according to Perozeni, et. al (Perozeni et al. 2019). Cells were frozen in liquid nitrogen after being dark adapted or after 60 minutes of illumination at 1500 μmol photons m^-2^ s^-1^ of red light. Fluorescence decay kinetics were then recorded by using Chronos BH ISS Photon Counting instrument with picosecond laser excitation at 447 nm operating at 50 MHz. Fluorescence emissions were recorded at 680 nm in with 4 nm bandwidth. Laser power was kept below 0.1μW.

---

**Figure 1––figure supplement 1.**
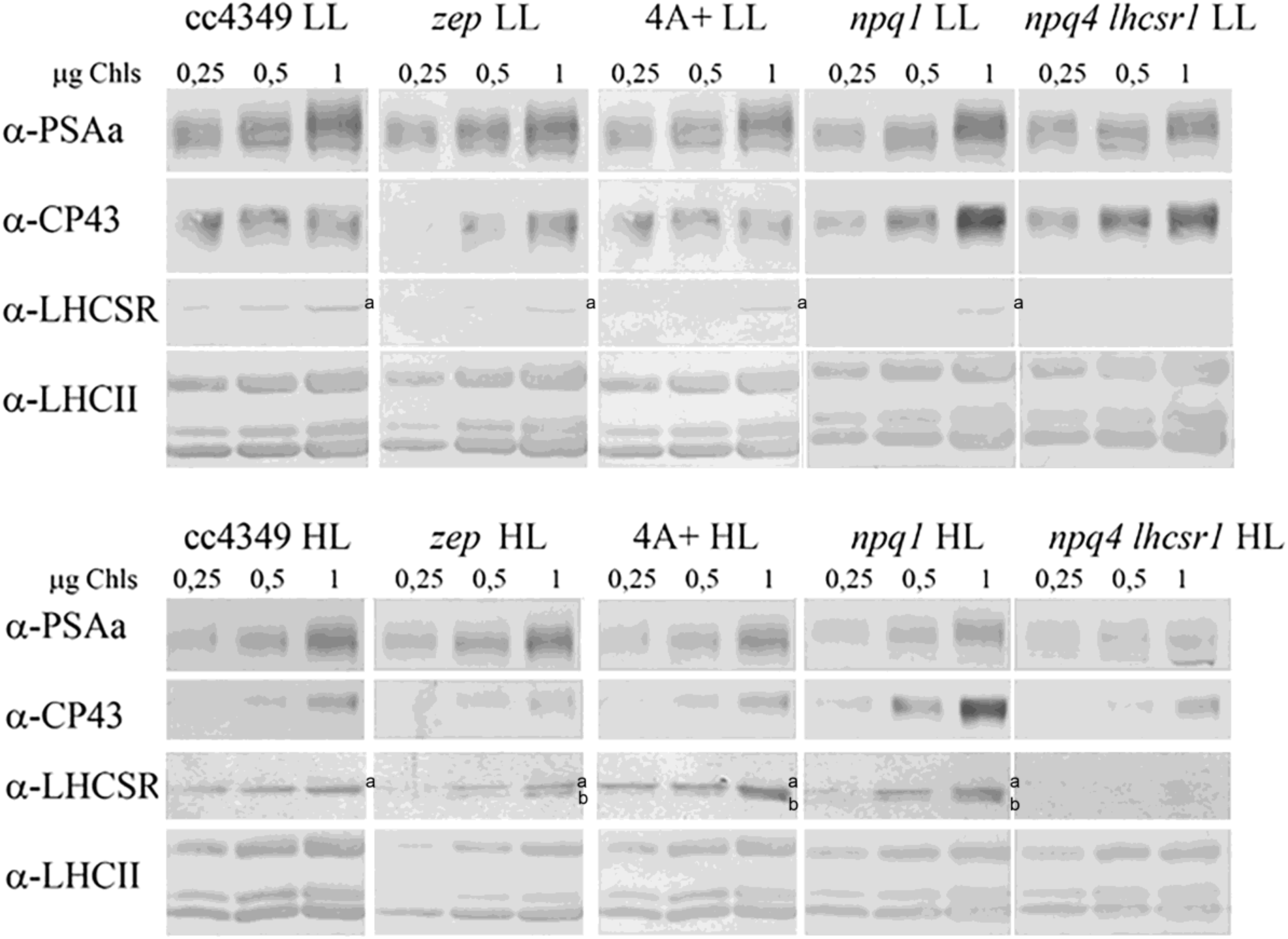
Immunoblot analysis of LHCSR accumulation in vivo. Total protein extracts from low Light (LL) or high light (HL) acclimated cells were loaded on a chlorophyll basis (μg of chlorophylls loaded are reported in the figure). Immunoblot analysis was performed using specific antibodys recogniziign PsaA, CP43, LHCSR3 (a) and LHCII subunits. The filter used for LHCSR3 detection was then used for LHCSR1 (b) detection using specific antibody. The results reported are representative of two independent experiments with different biological replicates and three technical replicates for each genotype in low light (LL) or high light (HL).

**Figure 1––figure supplement 2.**
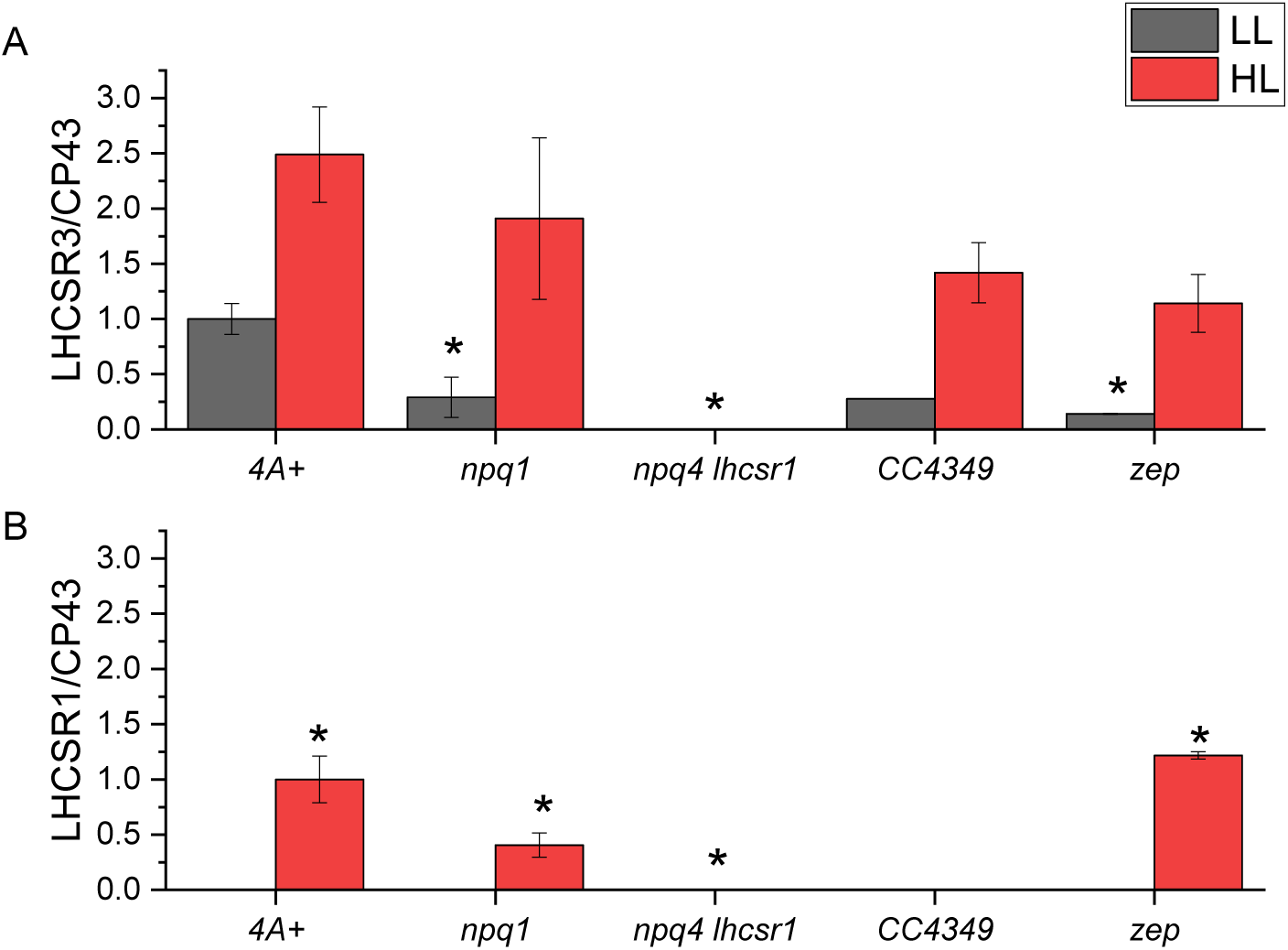
Quantification of LHCSR3 and LHCSR1 accumulation per PSII. Immunoblotting results were analyzed by densitometry. LHCSR3 (A) and LHCSR1 (B) content was then normalized to the amount of CP43 as a reference for PSII. The results obtained were then normalized to the 4A+ LL case in the case of LHCSR3 and 4A+ HL case in the case of LHCSR1. The results reported are representative of two independent experiments with different biological replicates and three technical replicates for each genotype in low light (LL) or high light (HL). Error bars are reported as standard deviation. The statistical significance of differences compared to WT (4A+ for *npq1* and *npq1 lhcsr1* mutants, CC4349 for *zep* mutant) is indicated as * (p<0.05), as determined by unpaired two sample t-test (N=3).

**Figure 1––figure supplement 3.**
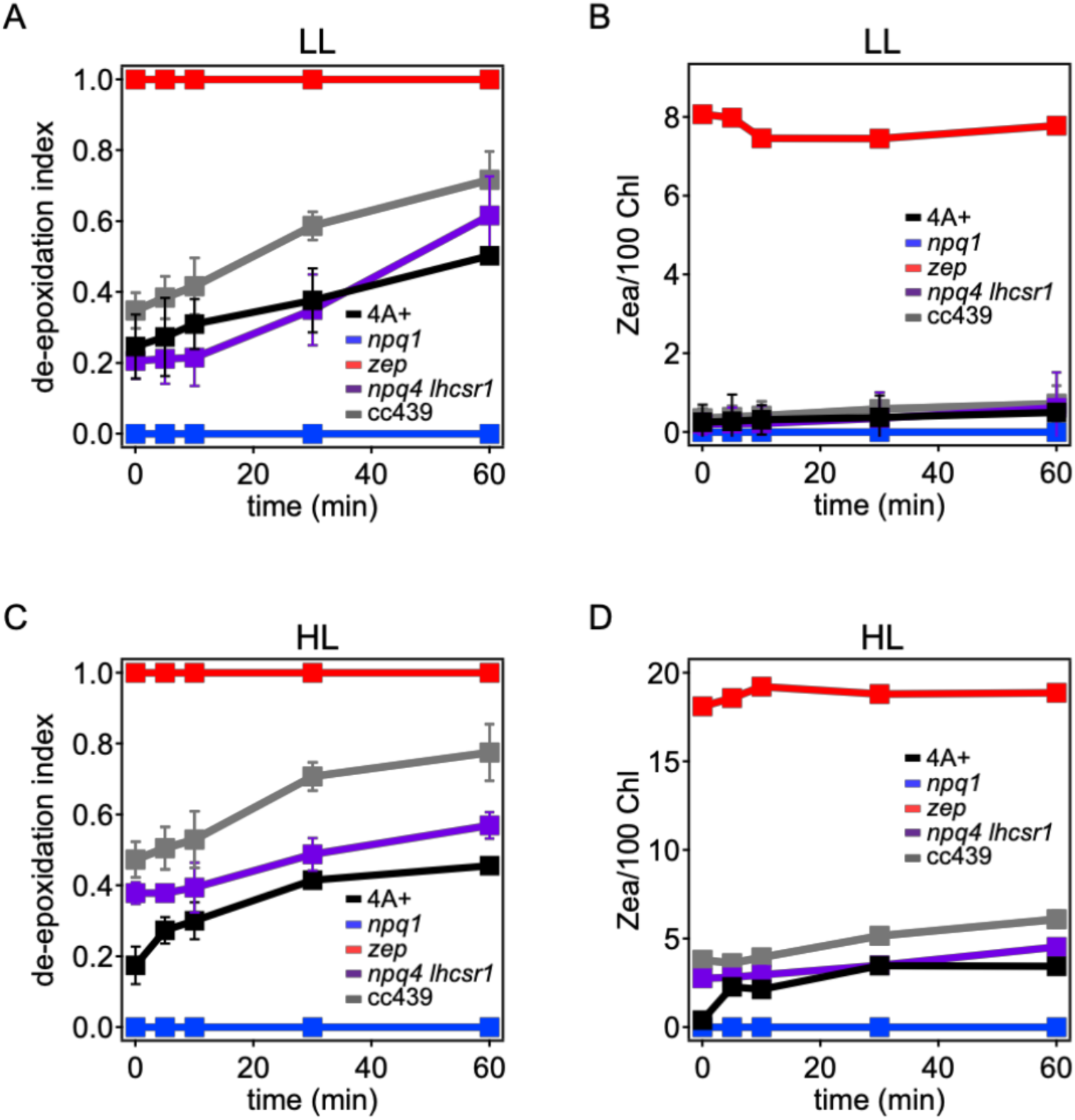
Violaxanthin de-epoxidation kinetics in *Chlamydomonas reinhardtii* WT and mutant strains. Violaxanthin de-epoxidation kinetics were measured upon 60 minutes of strong light treatment (red light 1500 μmol m-2s^-1^) of low light (LL) or high light (HL) acclimated cells. Panel A, C: de-epoxidation indexes at different time points. Panel B, D: zeaxanthin content per 100 chlorophylls. De-epoxidation index was calculated from the molar concentration of zeaxanthin, anteraxanthin and violaxanthin as ([zeaxanthin]+ [anteraxanthin]/2)/([zeaxanthin]+ [anteraxanthin]+ [violaxanthin]). The results reported are representative of three independent biological replicates for each genotype in LL or HL. Error bars are reported as standard deviation (N=3).

**Figure 1––figure supplement 4.**
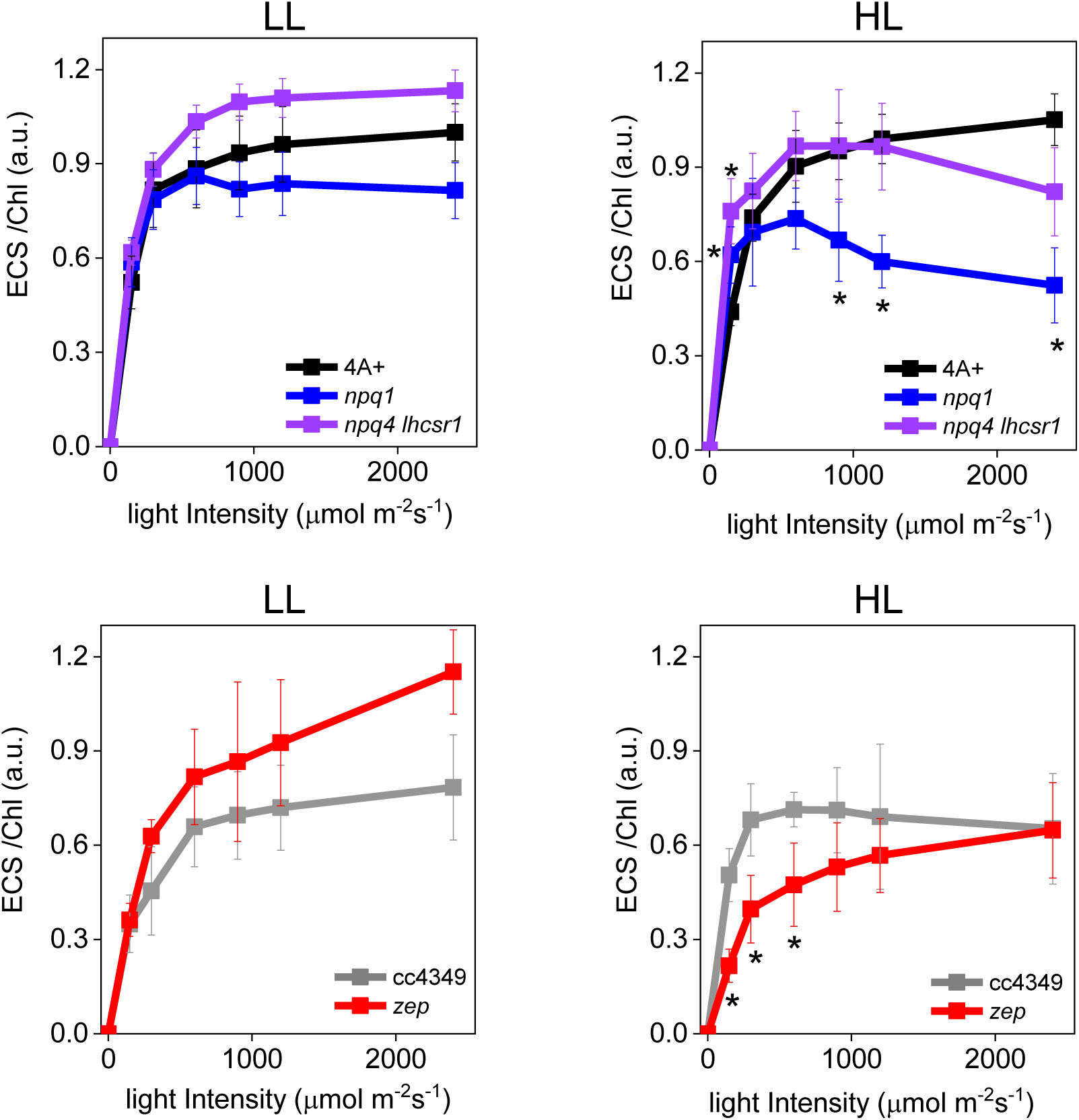
Electrochromic shift measurements at different light intensities in low light and high light acclimated *Chlamydomonas reinhardtii* cells. Electrochromic shift (ECS) measurements were performed in WT (4A+ and cc4349) and mutant strains (npq4 lhcsr1, *npq1* and *zep*) acclimated to low (Panel A and B) or high (Panel C, D) light. Genotypes having the same background are shown in the same Panel. The results reported are representative of three independent biological replicates for each genotype in LL or HL. Error bars are reported as standard deviation. The statistical significance of differences compared to WT (4A+ for *npq1* and *npq1 lhcsr1* mutants, CC4349 for *zep* mutant) is indicated as * (p<0.05), as determined by unpaired two sample t-test (N=3).

**Figure 1––figure supplement 5.**
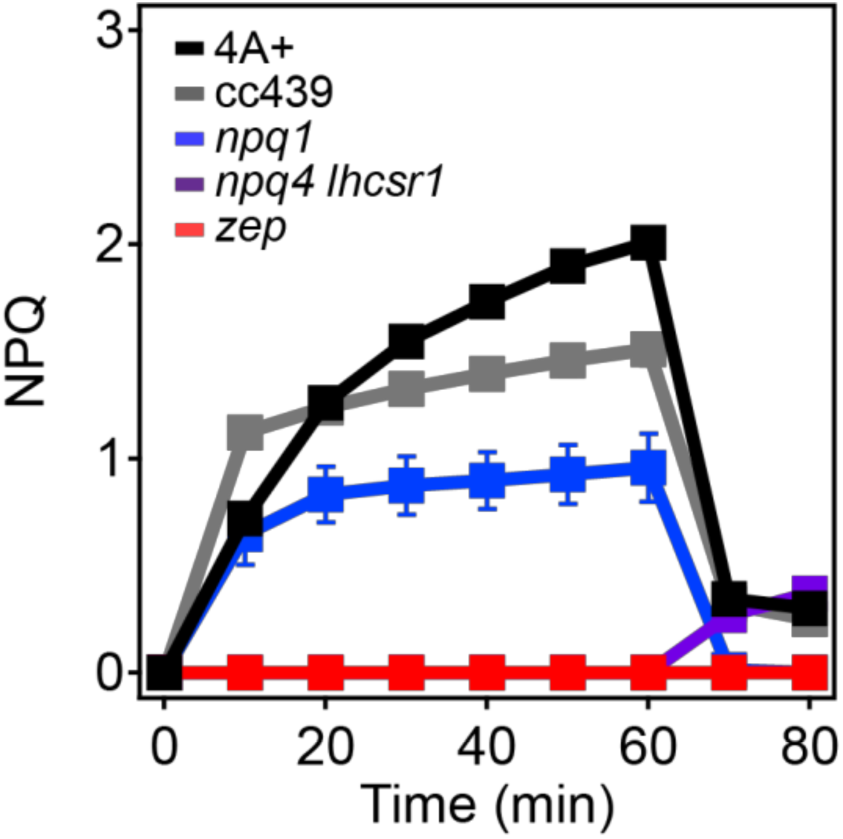
NPQ induction in low light acclimated Chlamydomonas reinhardtii cells. NPQ induction kinetics measured upon 60 minutes of high light (2000 μmol m-2s^-1^) followed by 20 minutes of dark recovery. The results reported are representative of three independent biological replicates for each genotype in LL or HL. Error bars are reported as standard deviation (n=3).

**Figure 1––figure supplement 6.**
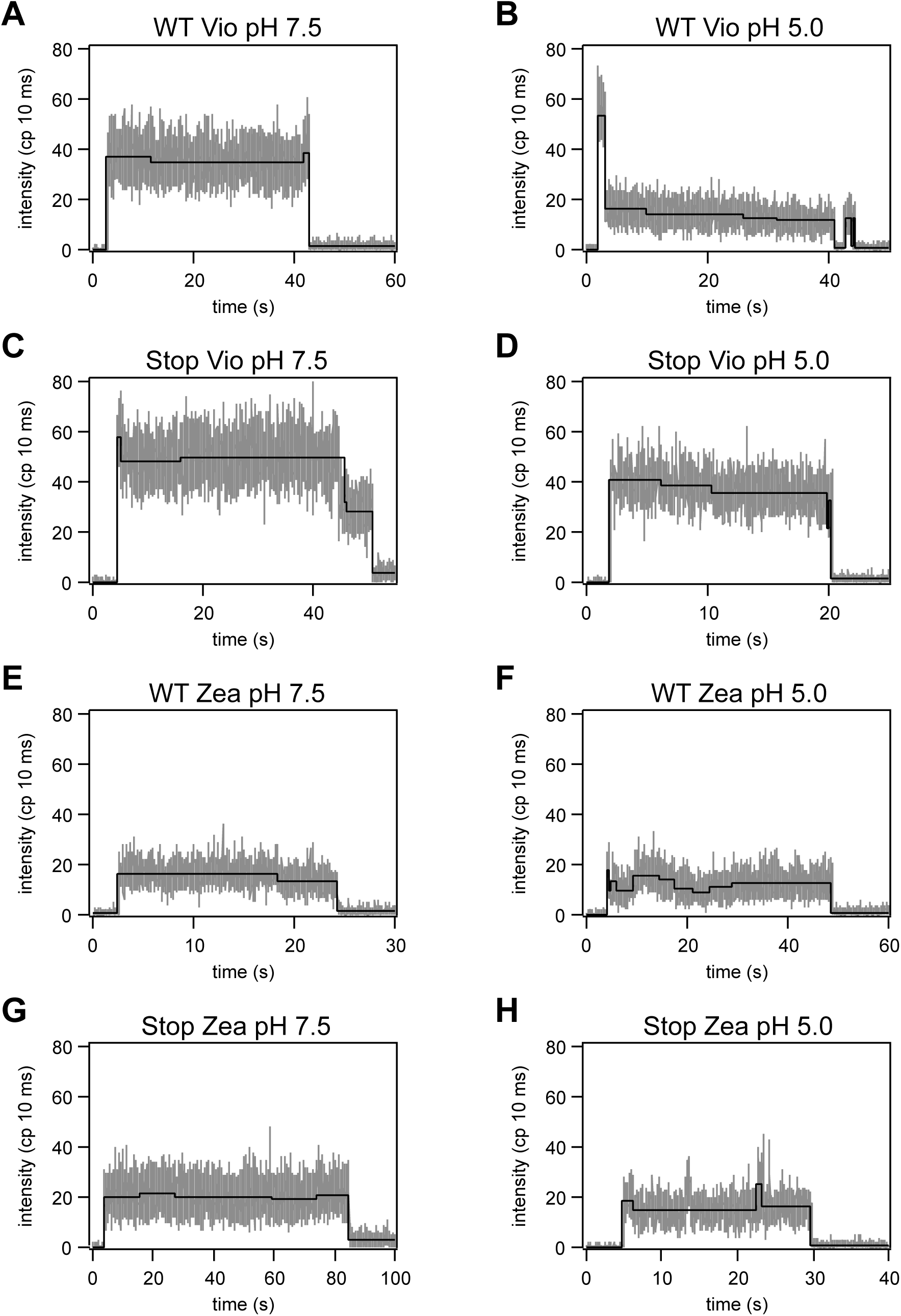
Representative fluorescence intensity traces. Fluorescence intensity traces for LHCSR3 complexes at pH 7.5 and 5.0. The intensity levels determined by the change-point-finding algorithm are shown in black.

**Figure 1––figure supplement 7.**
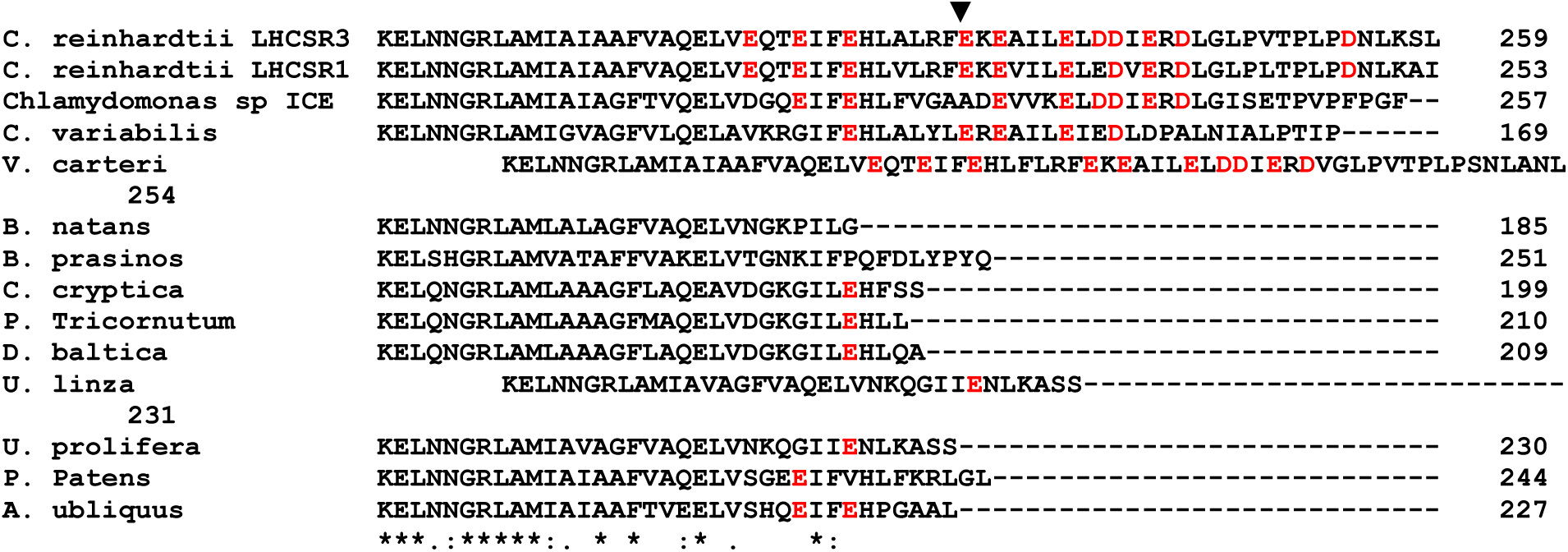
Alignment of LHCSR-like proteins: protonatable residues are red written while insertion site for TAA mutation to generate STOP mutant is indicated by black arrow. Protein number for LHCSR-like proteins used for alignment are listed below: XP_001696064.1 Chlamydomonas reinhardtii LHCSR3, XP_001696125.1 Chlamydomonas reinhardtii LHCSR1, XP_002948670.1 Volvox carteri f. nagariensis, ADP89594.1 Chlamydomonas sp. ICE-L LHCSR2, XP_001768071.1 Physcomitrella patens LHCSR2, ABD58893.1 Acutodesmus obliquus, ADY38581.1 Ulva linza, ADU04518.1 Ulva prolifera, XP_005848576.1 Chlorella variabilis, XP_002178699.1 Phaeodactylum tricornutum, AHH80644.1 Durinskia baltica, AA05890.1 Bigelowiella natans, CCO66741.1 Bathycoccus prasinos.

**Figure 1––figure supplement 8.**
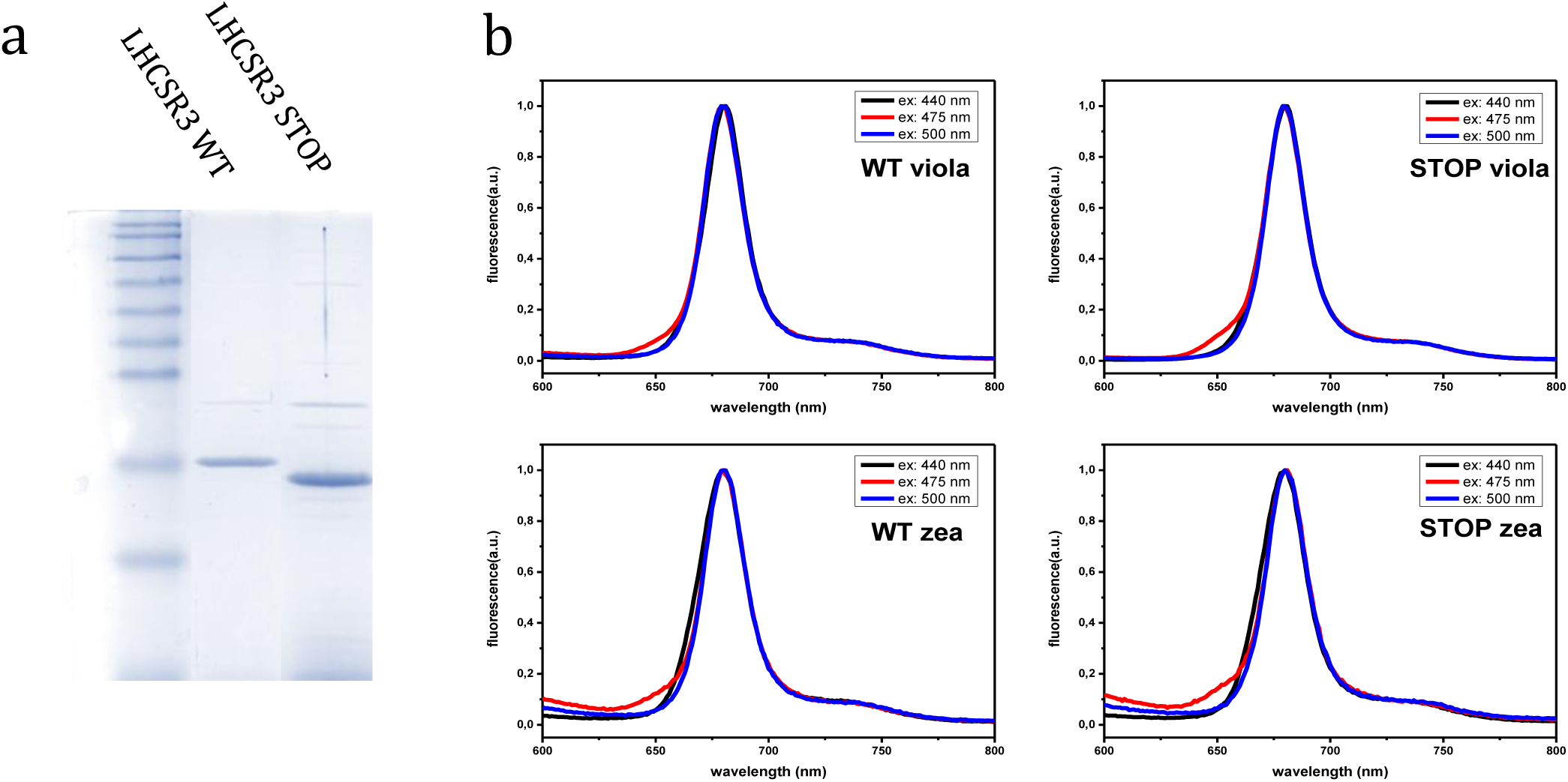
(a) Coomassie brilliant blue stained SDS-page of LHCSR3 apo-protein separated on Tris-Glycine 12%. LHCSR3 STOP protein shows high mobility conferred by its shorter C-terminal with respect to LHCSR3 WT; (b) Fluorescence spectra of LHCSR3 complexes measured at room temperature at different excitation wavelengths (reported in each panel). The results reported are representative of two independent experiments.

**Figure 1––figure supplement 9.**
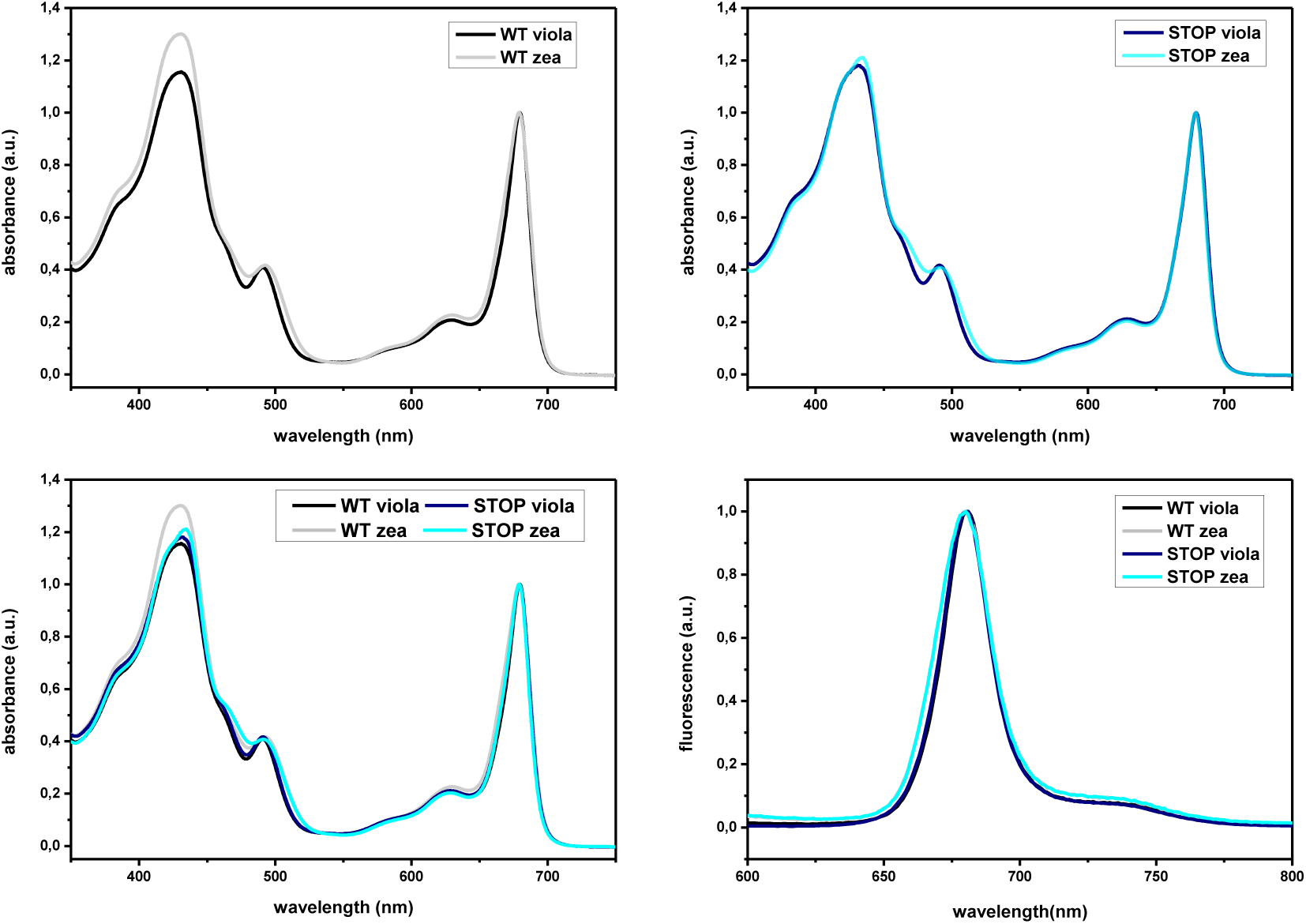
Absorption and fluorescence emission spectra of LHCSR3 WT and STOP. (a-c) Absorption spectra of WT (a) or STOP (b) refolded with violaxanthin or zeaxanthin. (d) Fluorescence emission spectra of LHCSR3 WT and STOP mutant upon excitation at 440 nm with both violaxanthin and zeaxanthin pigments composition. The results reported are representative of two independent experiments.

**Figure 2––figure supplement 1.**
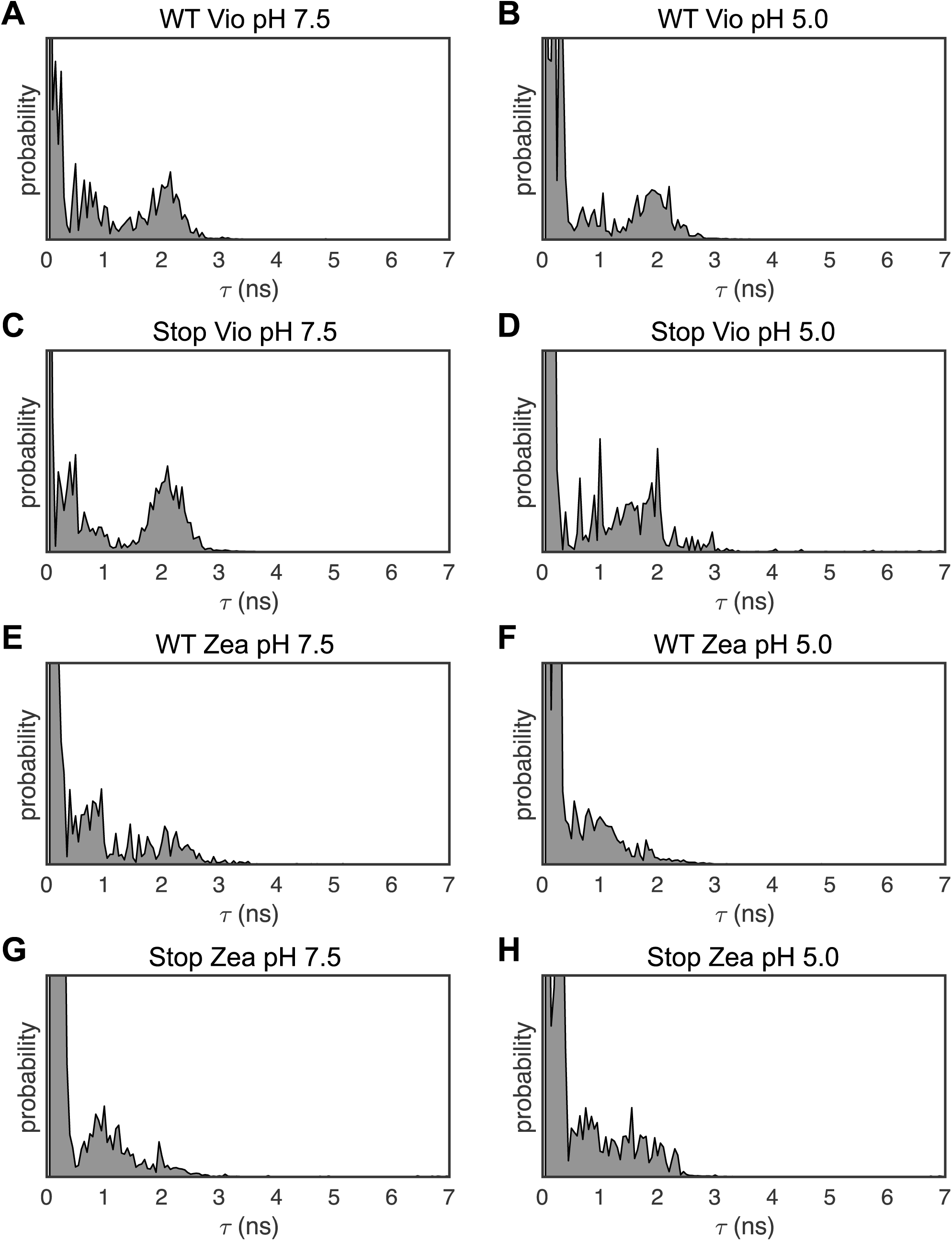
1D Lifetime distributions. Lifetime distributions for LHCSR3 determined using a one-dimensional inverse Laplace transform (1D-ILT) of the 1D fluorescence lifetime decay. Lifetime states identified from the 1D distribution were used as initial parameters in the fit to the 2D distributions.

**Figure 2––figure supplement 2.**
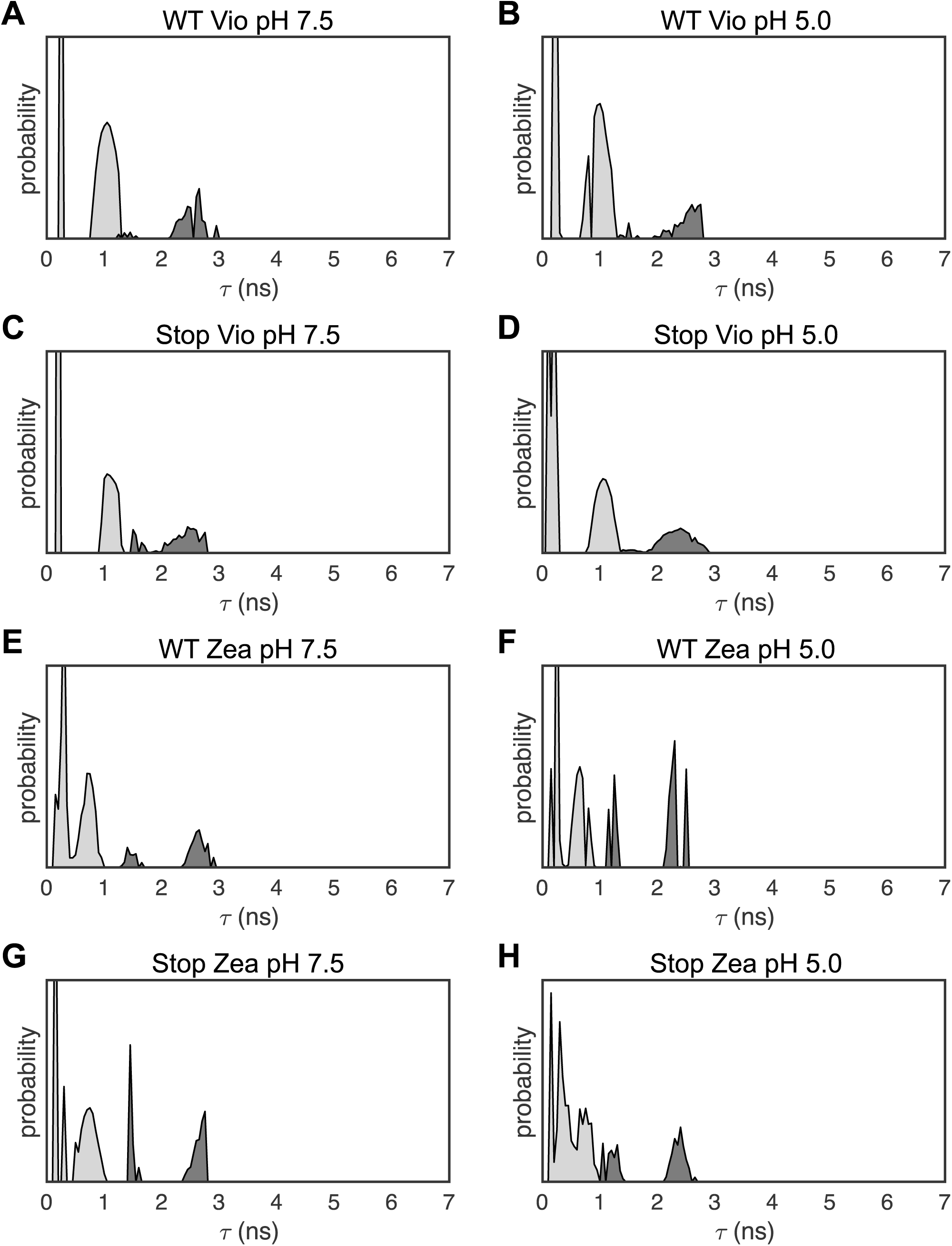
2D Lifetime distributions. Lifetime distributions for LHCSR3 complexes at pH 7.5 and 5.0 determined using the maximum entropy method (MEM) to perform a 2D-ILT.

**Figure 2––figure supplement 3.**
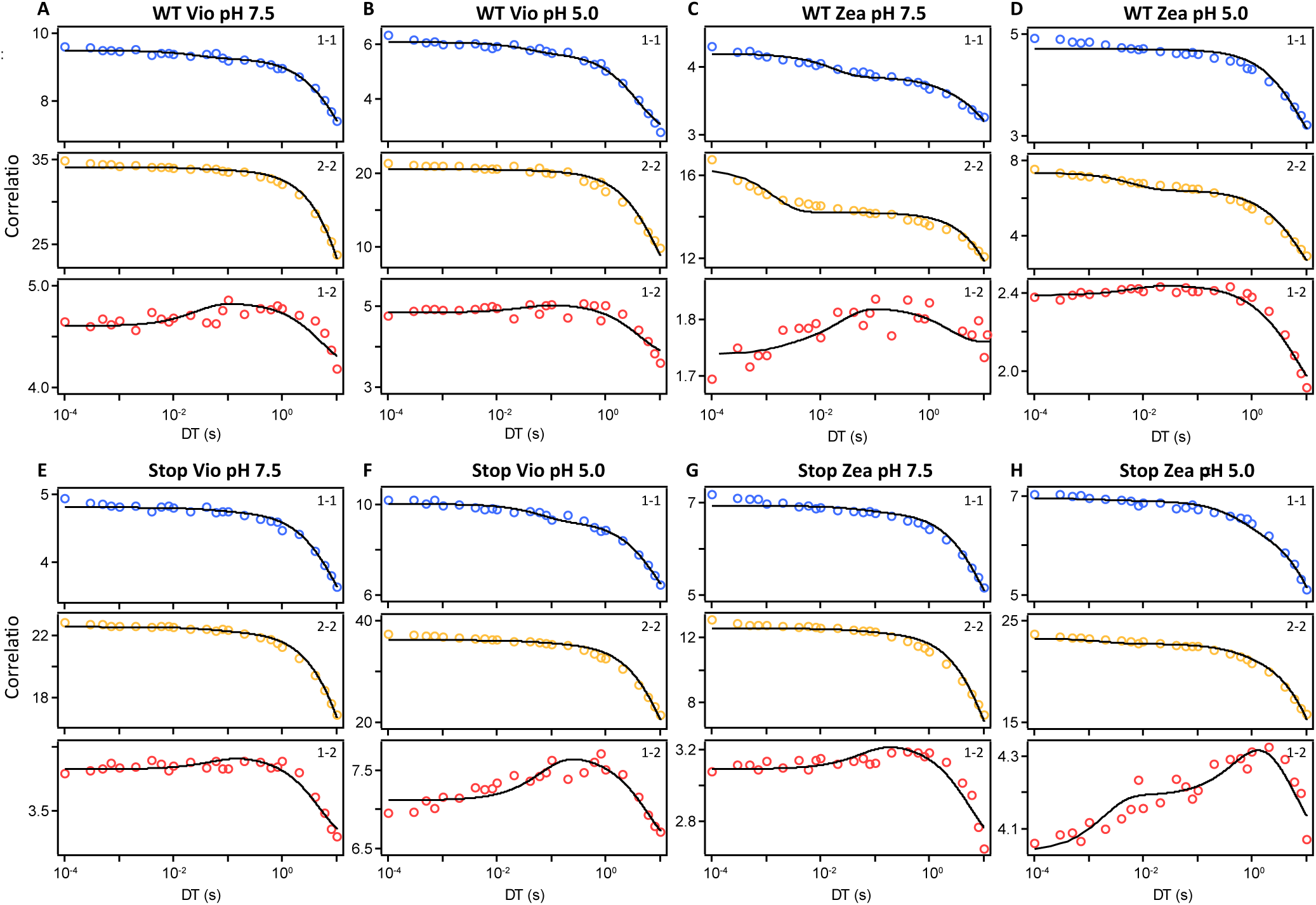
Correlation analysis of LHCSR3 complexes. Correlation function estimated from the 2D-FLC analysis of single LHCSR3 complexes with Vio at pH 7.5 (A) and pH 5.0 (B), Zea at pH 7.5 (C) and 5.0 (D), stop mutants with Vio at pH 7.5 (E) and pH 5.0 (F) and stop mutants with Zea at pH 7.5 (G) and pH 5.0 (H). The correlation curves for auto (1-1 and 2-2) and cross correlations (1-2) are shown in blue, yellow and red, respectively. The black line shows the fitting curve calculated using the model function given by equation described in the Methods.

**Figure 2––table supplement 1.**
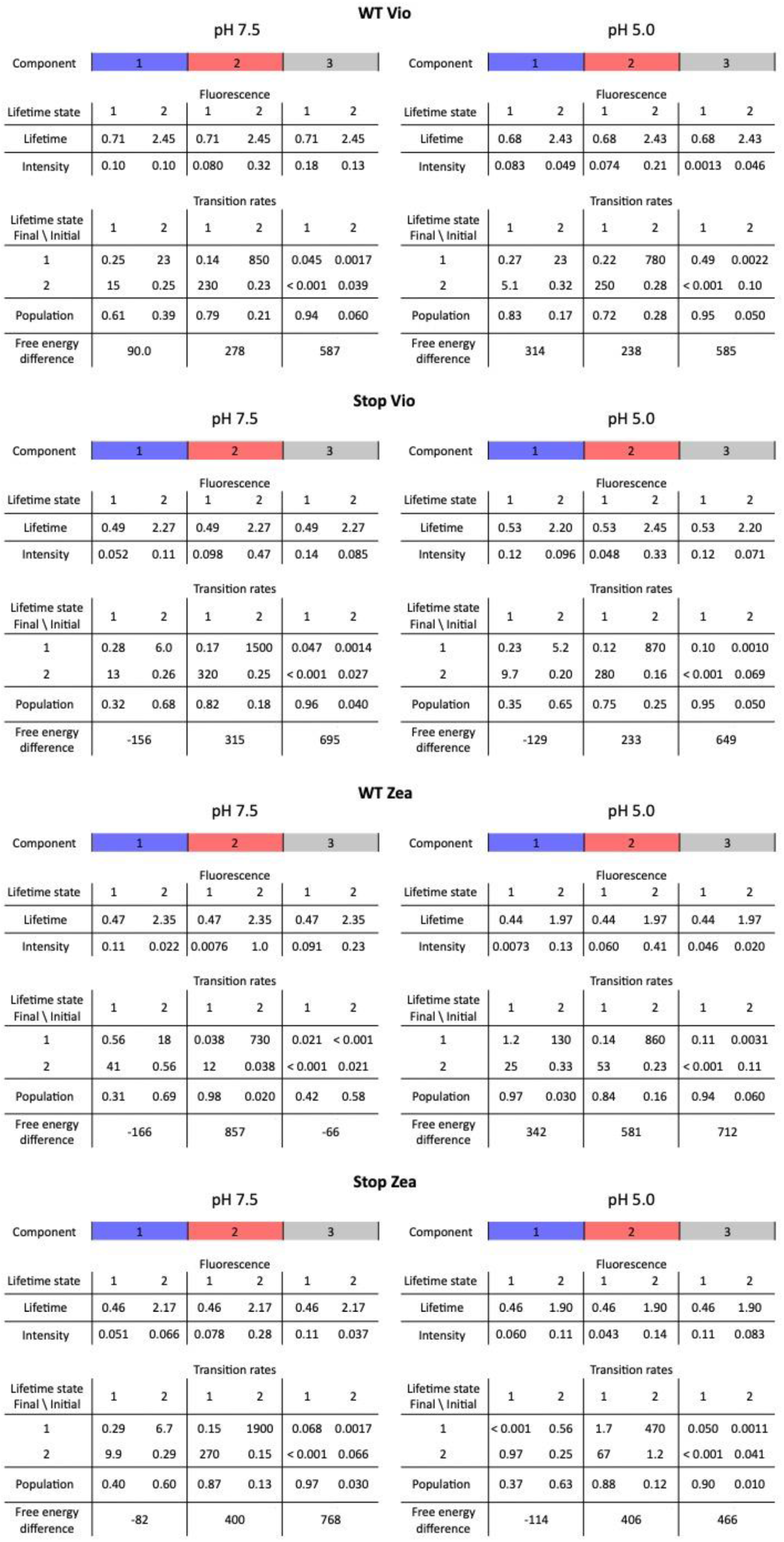
Summary of dynamic properties estimated by the correlation fitting analysis. The fluorescence intensity and population of the initial and final state and the transition rates between these states for each component (blue, red, gray) were determined by global fitting of the correlation functions shown in Figure S18. The free-energy differences were given by the equation described in the Methods. The fluorescence intensity is a relative intensity that is normalized by the total measurement time for each sample and by a scaling factor to set the maximum intensity to be 1.

**Figure 2––table supplement 2.** Pigment binding properties of recombinant LHCSR3 WT and STOP refolded in vitro. Binding pigments are reported referred to 7 Chlorophyll. The results reported are representative of two independent experiments. Errors are reported as standard deviation.

|  | Total<br>Chlorophylls | Chlorophyll a | Chlorophyll b | Chla/Chlb | Neoxanthin | Violaxanthin | Anteraxanthin | Lutein | Zeaxanthin | Total<br>Carotenoids | Chl/Car |
| --- | --- | --- | --- | --- | --- | --- | --- | --- | --- | --- | --- |
| WT VIOLA | 7 | 6,59 ± 0,00 | 0,41 ± 0,00 | 16,16 ± 0,16 | 0,00 ± 0,00 | 0,98 ± 0,00 | 0,09 ± 0,03 | 1,02 ± 0,02 | 0,08 ± 0,01 | 2,17 ± 0,02 | 3,22 ± 0,02 |
| STOP VIOLA | 7 | 6,38 ± 0,01 | 0,62 ± 0,01 | 10,33 ± 0,12 | 0,00 ± 0,00 | 0,99 ± 0,07 | 0,07 ± 0,02 | 0,95 ± 0,08 | 0,03 ± 0,04 | 2,04 ± 0,05 | 3,43 ± 0,08 |
| WT ZEA | 7 | 6,82 ± 0,01 | 0,18 ± 0,01 | 37,58 ± 2,82 | 0,00 ± 0,00 | 0,27 ± 0,00 | 0,00 ± 0,00 | 0,31 ± 0,04 | 1,41 ± 0,02 | 1,99 ± 0,03 | 3,51 ± 0,05 |
| STOP ZEA | 7 | 6,81 ± 0,00 | 0,19 ± 0,00 | 36,56 ± 0,35 | 0,00 ± 0,00 | 0,22 ± 0,00 | 0,00 ± 0,00 | 0,44 ± 0,47 | 1,09 ± 0,02 | 1,75 ± 0,00 | 3,99 ± 0,00 |

**Figure 3––figure supplement 1.**
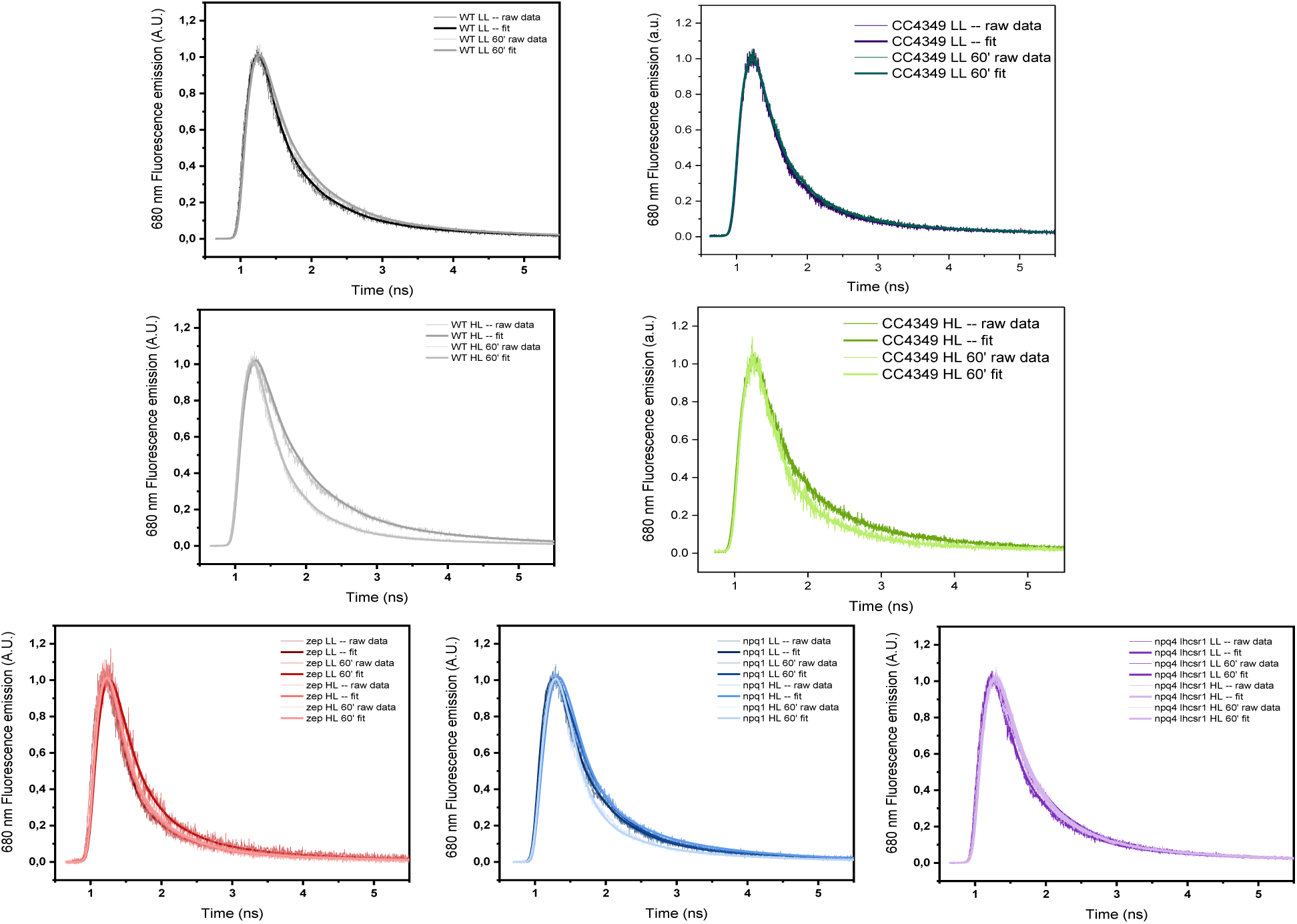
77K raw and fitted traces acquired by TCSPC of *Chlamydomonas reinhardtii* WT (4a+) and mutant strains. The results reported are representative of two independent experiments with two independent biologic replicates.

**Figure 3––figure supplement 2.**
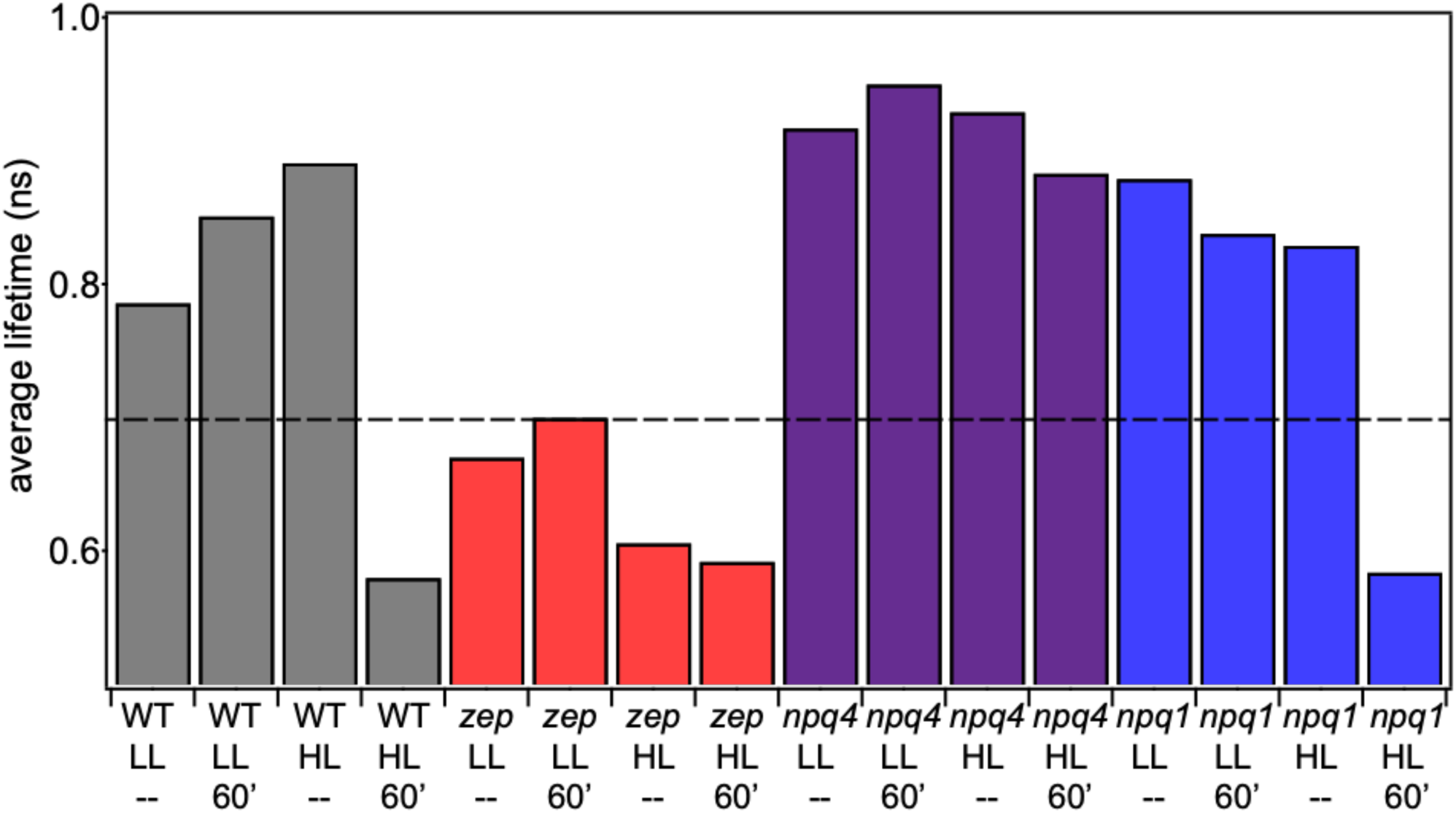
Average fluorescence lifetime for WT (4A+) and mutant strains under all light conditions. The other WT strain (cc4349) has similar values. Above dotted line is considered unquenched and below dotted line is considered quenched. The results reported are representative of two independent experiments with two independent biologic replicates.

**Figure 3––table supplement 1.**
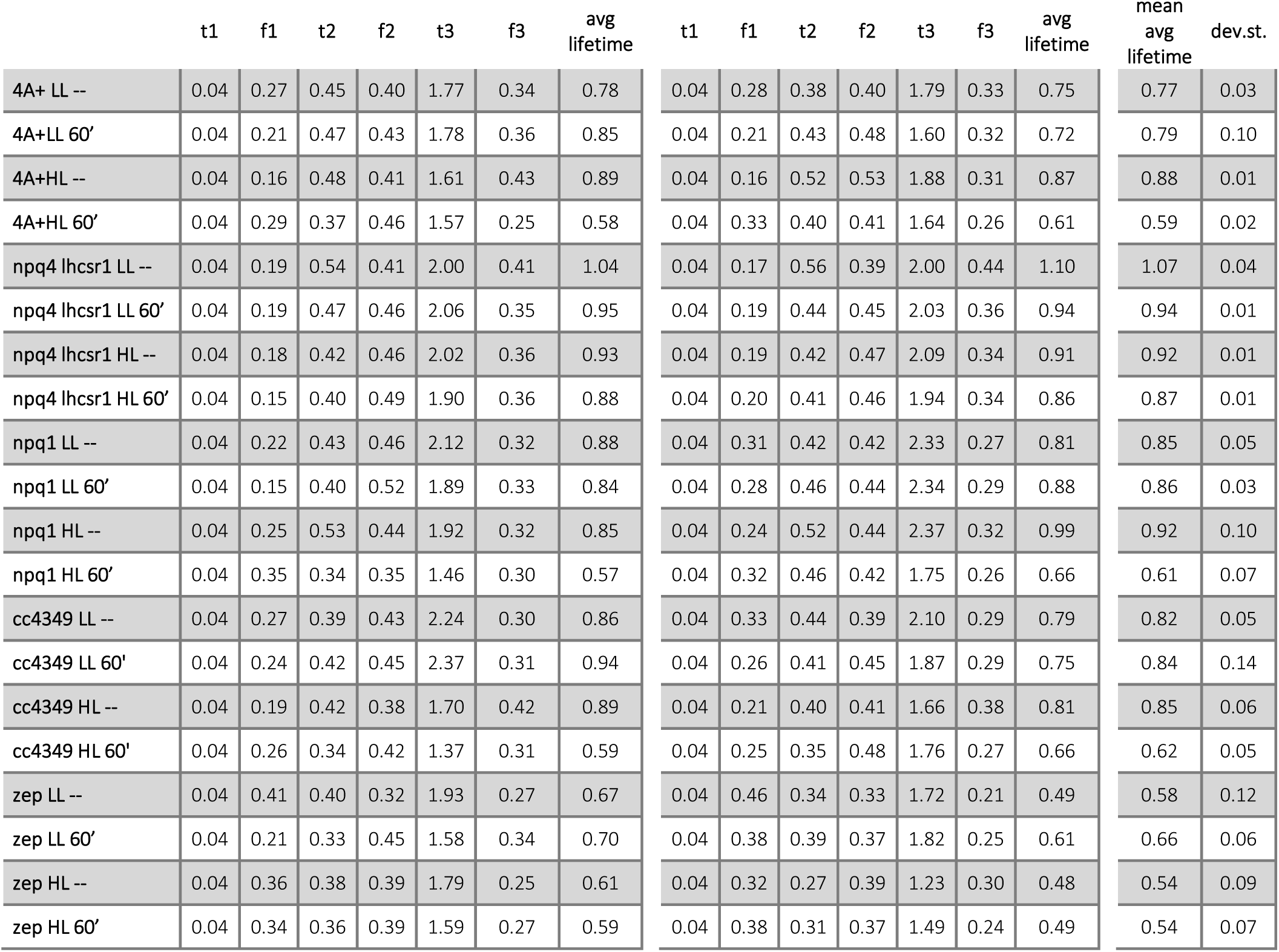
77K time resolved fluorescence analysis and average fluorescence decay lifetimes of whole cells. Kinetics were fitted with a three-exponential decay function using Vinci 2 software from ISS. Fractions (fi) and time constants (τi) are reported. Average fluorescence lifetimes were calculated as Σfiτi. The results reported are representative of two independent experiments with two independent biologic replicates.

**Figure 3––figure supplement 3.**
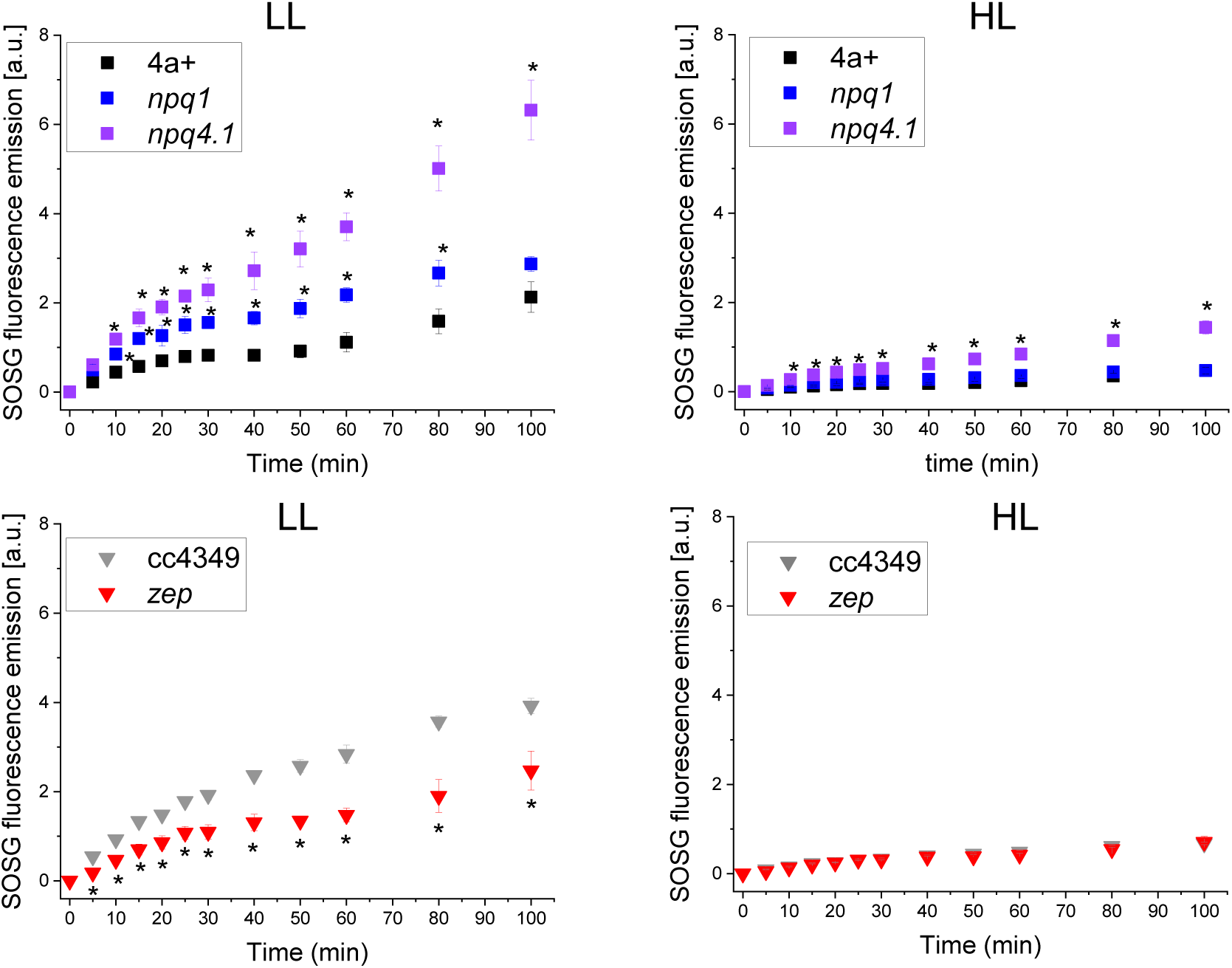
Kinetics of singlet oxygen production in WT and mutant strains. Singlet oxygen production rate was measured by the singlet oxygen sensor green (SOSG) fluorescence probe. Low light (LL) or high light (HL acclimated samples were exposed to strong red light (2000 μmol m^-2^s^-1^) and singlet oxygen production rate was probed at the different time points by following SOSG fluorescence at 530 nm. Genotypes having the same background are shown in the same Panel. The results reported are representative of three independent biological replicates for each genotype in LL or HL. Error bars are reported as standard deviation (n=3). The statistical significance of differences compared to WT (4A+ for *npq1* and *npq1 lhcsr1* mutants, CC4349 for *zep* mutant) is indicated as * (p<0.05), as determined by unpaired two sample t-test (N=3).

**Figure 3––figure supplement 4.**
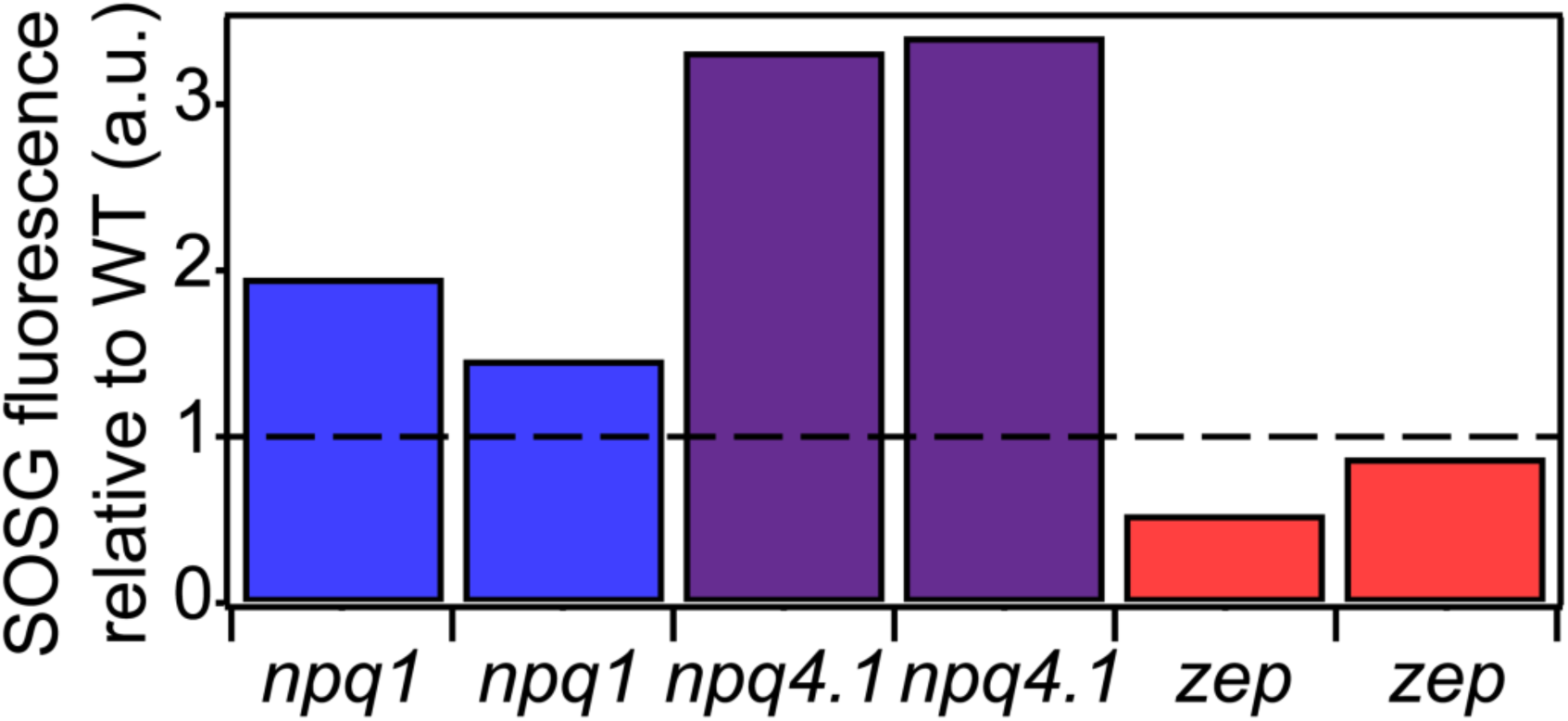
Singlet oxygen production in WT and mutant strains. Singlet oxygen production rate was measured by the singlet oxygen sensor green (SOSG) fluorescence probe. Low light (LL), left, or high light (HL), right, acclimated samples were exposed to strong red light (2000 μmol m^-2^s^-1^), right. Singlet oxygen production rates relative to WT (4A+ for *npq1* and *npq4 lhcsr1*, cc4349 for *zep*). The results reported are representative of three independent biological replicates for each genotype in LL or HL. Unnormalized data are reported in figure supplement 3.

